# Balancing competing effects of tissue growth and cytoskeletal regulation during *Drosophila* wing disc development

**DOI:** 10.1101/2022.09.28.509971

**Authors:** Nilay Kumar, Kevin Tsai, Mayesha Sahir Mim, Jennifer Rangel Ambriz, Weitao Chen, Jeremiah J. Zartman, Mark Alber

## Abstract

Cytoskeletal structure and force generation within cells must be carefully regulated as the developing organ grows to reach a final size and shape. However, how the complex regulation of multiple features of tissue architecture is simultaneously coordinated remains poorly understood. Through iterations between experiments and novel computational multi-scale model simulations, we investigate the combined regulation of cytoskeletal regulation and proliferation in the growing wing imaginal disc. First, we found through experiments and calibrated model simulations that the local curvature and nuclear positioning of cells in the growing wing disc are defined by patterning of nested spatial domains of peaks in apical and basal contractility. Additionally, predictions from model simulations that incorporate a mechanistic description of interkinetic nuclear migration demonstrate that cell proliferation increases the local basal curvature of the wing disc. This is confirmed experimentally as basal curvature increases when growth and proliferation are increased through insulin signaling. In surprising contrast, we experimentally found that Decapentaplegic (Dpp), the key morphogen involved in both growth control and patterning of the anterior-posterior axis, counteracts increases in tissue bending due to cell proliferation via a combined mechanism that balances the competing impacts of both proliferation and patterning of cell contractility. Overall, the high conservation of these regulatory interactions suggests an important balancing mechanism through dual regulation of proliferation and cytoskeleton to meet the multiple criteria defining tissue morphogenesis.

## Introduction

The final shape of an organ is a result of the dynamic interplay between several cell-level processes^1–4^. Progress in uncovering the regulation of organ shape control can be achieved by using systems-level, multi-scale approaches^5^. A major challenge in reverse-engineering biological systems arises from the inherent complexity of non-linear interactions between proteins and cells that occur over multiple spatial and temporal scales^6^. In particular, the regulation of features such as cell height, local tissue curvature, and nuclear positioning is critical for controlling the morphogenesis of multicellular organisms^7–9^. Untangling the regulation of multiple subcellular features that compete with each other over both short and long time scales requires systematic analysis that incorporates highly complex biologically calibrated computational model simulations coupled with quantitative experimental analysis^10,11^.

Of note, epithelial tissues act as external protective lining for internal organs^12^. Extreme topological changes in the form of “folding” and “flattening” that occur within epithelial tissues during organ development are necessary for the manifestation of its 3-D structure^13^ and functions^14^. The structural changes occurring within a developing organ are mediated through individual cell shape changes arising due to a spatiotemporal patterning of cytoskeletal regulators. In the case of *Drosophila melanogaster*, a fundamental question lies in determining the systems-level mechanism driving the formation and maintenance of wing disc shape during the larval stage and the connection between such shape acquisition with patterning of cell identities and cellular properties including growth^13–15^. In particular, proper shape acquisition plays a central role in the ability of a tissue or organ to function and contribute to the fitness of the organism. For instance, the final wing shape is a key determinant in flight performance^16^.

In this work, we developed a systems-level understanding of the independent roles of dynamics in cytoskeletal regulation and cell proliferation in the shaping of an organ using the *Drosophila* wing imaginal disc as a biological model. Nuclear positioning and local basal curvature are key attributes defining cell and tissue architecture. Recent studies suggest that tissue architecture contributes to cell proliferation, but the mechanisms of such cross-talk are generally poorly understood^17^. Previous studies in the wing imaginal disc have shown that an increase in apical curvature promotes the apical positioning of nuclei in a nuclear density-independent manner to promote proliferation^8^. Here, we describe the role of cell proliferation in regulating the tissue basal curvature through a combination of experiments and simulations using a novel multiscale model that we developed. The experimentally calibrated computational framework opens up new avenues for the exploration of a feedback loop present between tissue shape and cell proliferation. In addition, we also show that cell proliferation alone cannot generate the correct shape without the actomyosin patterning. For the first time, we have introduced a cell division algorithm that captures the interkinetic nuclear migration and anisotropic mitotic rounding process, allowing us to closely replicate experimental observations of subcellular dynamics^8,9^.

We further show that a key requirement for the wing disc to acquire its experimentally observed shape is a balance in the form of a tug-of-war^18^ between the contractile forces originating from the cooperative activity of actin filaments and the myosin motors manifested both apically and basally within each cell. Improper patterning of the actin and myosin (actomyosin) activities can lead to the wing disc acquiring shapes that qualitatively agree with mutant wing disc shapes. Both the patterning and the actual contractile force generated also influence the positioning of nuclei within each cell, which is found to follow a specific pattern in the wild-type wing disc.

During *Drosophila* wing development, Decapentaplegic (Dpp), a key morphogen, activates and patterns the Rho family of small GTPases to generate a gradient in cell height. Rho is involved in the phosphorylation of non-muscle myosin II^19^, which pulls on actin to generate contractile forces to drive folding within the tissue^20^. Theoretical studies performed on embryonic epithelia also reveal that epithelial sheet thickness is primarily determined through mechanical interactions between microfilaments, microtubules, and cell adhesion systems^21^. Adhesion proteins like cadherins and Integrins are also critical for holding the tissue together as it grows in size. For instance, Integrins adhere the basal surfaces of cells to the extracellular matrix (ECM)^22^. Integrins play multiple roles during development^23^, such as acting as a cell surface receptor involved in signal transduction into the cell through the ECM^24^. Clustering of Integrins is also known to facilitate the formation of focal adhesion complexes facilitating cell migration and division reported across multiple systems^25^. A recent study also highlights the importance of E-Cadherin, a second class of adhesion molecules, in maintaining coordination in cell-cell communication and self-organization during embryonic development^26^.

Regulation of cell mechanics within the growing pseudostratified epithelium is critical for its shape maintenance. Exponential organ growth is accompanied by a significant increase in cell biomass. It should be also noted that the timescale of seconds to minutes in cytoskeletal regulation varies from that of hours in cell proliferation. The difference in timescales also makes it difficult to understand how an organ shapes during high proliferative phases. A recent study by Tozluoǧlu et al. shows the role of planar differential growth rates in determining fold position along the dorsal-ventral axis of the wing imaginal disc^27^. Some work has also been done to explore this regulation in the other direction, i.e., how the shape of an organ influences proliferation^28,29.^ However, how proliferation locally shapes an organ during development is still poorly understood.

A key model system for studying the regulation of morphogenesis across spatiotemporal scales and regulatory modes is the wing disc of the fruit fly. The wing imaginal disc at the 3^rd^ instar larval stage consists of the pouch, hinge, and notum. The pouch cells mature to form the adult wing of a fly (Fig. 1A). Most models for growth in *Drosophila* wing imaginal disc define two axes of development, which are perpendicular to each other. The development in directions perpendicular to these axes is governed by the morphogens such as Wingless and Dpp (Fig. 2A-iii)^30^. Wing imaginal discs resemble a dome shape with an anisotropy in patterning of basal curvature that changes with its age. At the early 3^rd^ instar larval stage, the wing imaginal disc is basally more curved at the anterior and posterior (lateral) ends to form bends as compared to the relatively flat medial region (Fig. 1B-i-v).

**Figure 1:**
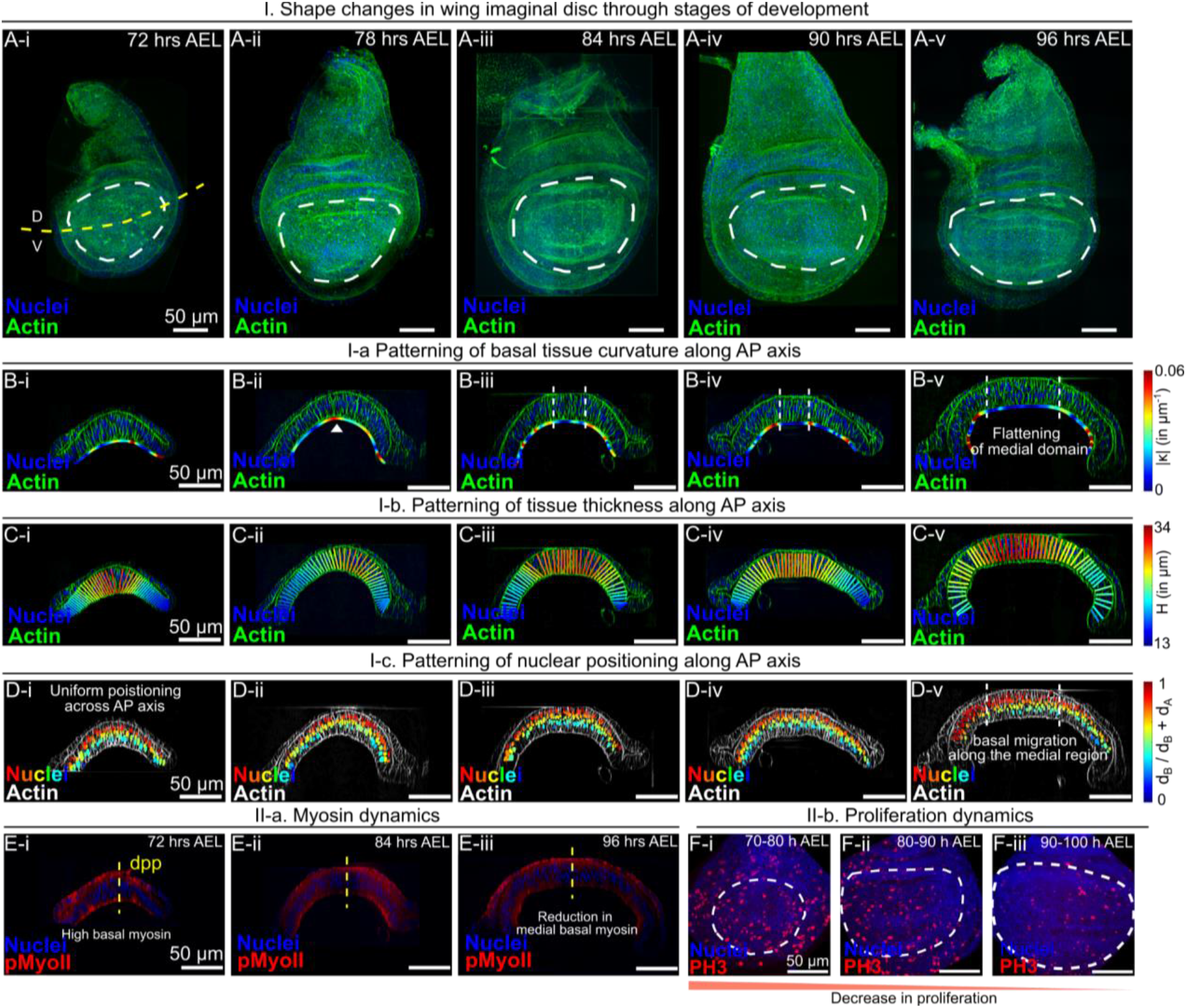
The central domain of the wing disc pouch flattens and thickens as it grows. **(A i-v)** Apical view of a wild type Ore-R wing imaginal disc at five different larval stages. Fluorescence signals in green and blue denote the patterning of actin and nuclei. **(B i-v)** Cross-sectional view of wing imaginal discs at five different developmental stages. The basal surface is overlaid with its local curvature. **(C i-v)** The AP cross-section of the wing imaginal disc is overlaid with color-coded lines representing the local tissue thickness profile in terms of height, H. **(D i–v)** AP cross-section of the wing imaginal discs is overlaid with color-coded nuclei representing the ratio of the distance of nuclei from basal surface (dB) to the sum of distances from apical (dA) and basal surfaces respectively. **(E i-iii)** AP cross-sections highlighting localization of pMyoII at different developmental stages. **(F i-iii)** Apical view of pouch showing dividing cells (PH3) at different developmental stages.

**Figure 2:**
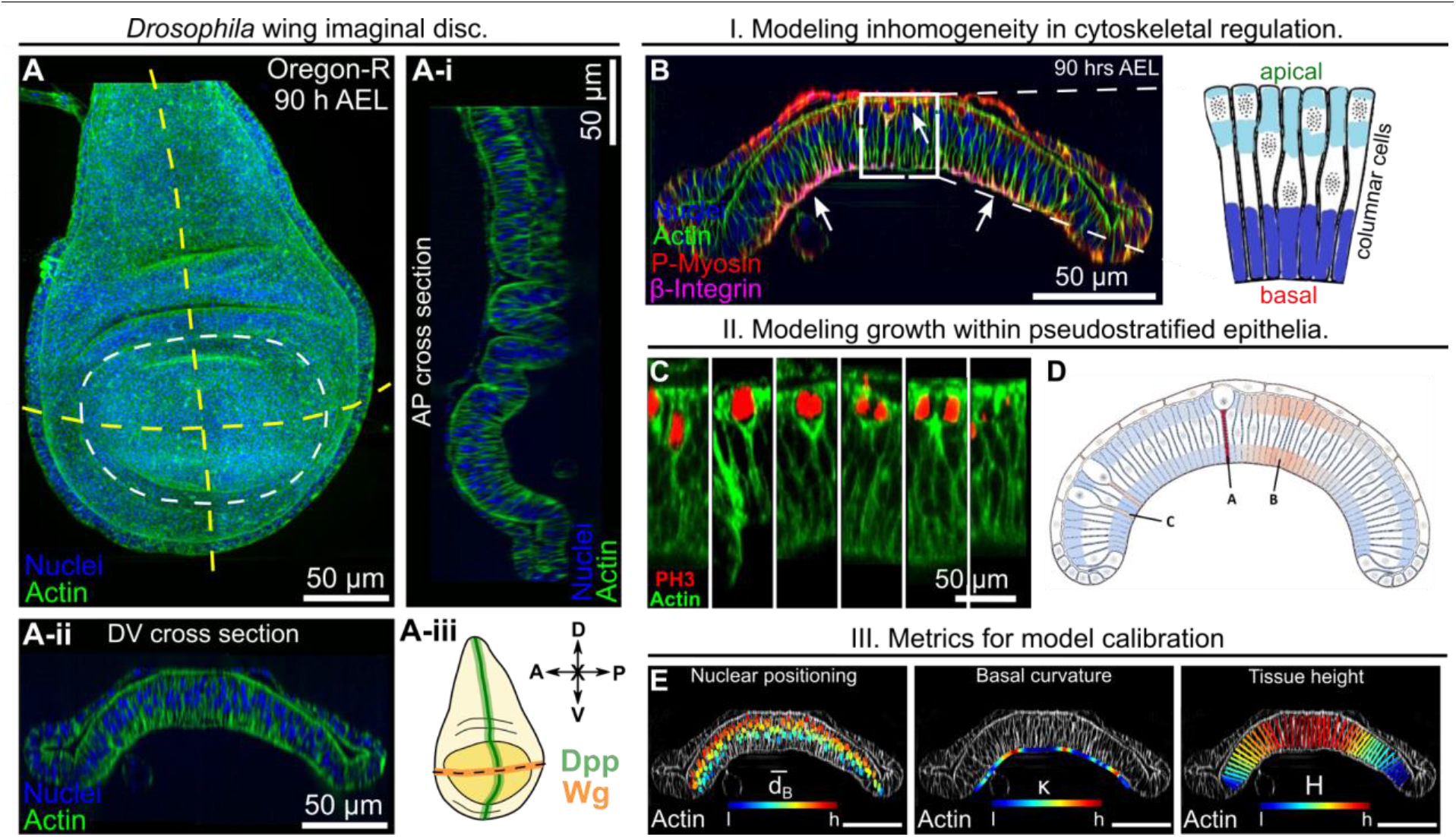
Multiscalar subcellular element simulation of proliferation in a pseudostratified epithelium predicts proliferation dynamics and cytoskeletal regulation. **(A)** Maximum intensity z plane projection showing the apical view of a *Drosophila* wing imaginal disc dissected from a larva at 90 hours AEL. **(A-i - Aii)** Cross-sectional views of the wing imaginal disc running parallel along the Anterior-Posterior (AP) and Dorsal-Ventral (DV) boundaries, respectively. **(A-iii)** A schematic illustrating patterning of the key morphogens Decapentaplegic (Dpp) and Wingless (Wg). The inset on top specifies compartment orientation for data visualization and analysis. **(B)** Apical and basal membrane contractility inside individual cells is modeled as linear springs connecting the corresponding membrane nodes. The number of contractile springs depends on the local actin intensity measured from experimental images. The spring coefficient for the contractile springs is assumed to be proportional to local myosin intensity. Parameters are determined based on experimental quantification. **(C)** Stages of interkinetic nuclear migration and cell division in wing disc cells. **(D)** Computational snapshot of simulated cell division. **(E)** Geometrical features defining the global tissue architecture and used as the metrics for calibration of the subcellular element model include: 1. Nuclear distance from the basal surface, dB, 2. Local basal curvature, κ and 3. Tissue thickness, H. Color-code represents lower (l) to higher (h) numerical quantity.

In this paper, we decipher the regulated balance of actomyosin-mediated contractility and cell proliferation in the shaping of an organ. We experimentally quantified the shape changes along the anterior-posterior (AP) section of the pouch during the late stages of *Drosophila* wing development. In particular, we observed an increase in tissue thickness, a decrease in local curvature of the basal surface and a shift in the nuclear positioning of nuclei toward the basal sice within the mid-section of the pouch as it grows to its final size (Fig. 1). To test whether the proposed mechanism of spatial patterning of cell mechanical properties, such as actomyosin contractility, leads to the emerging shape observed in experiments, we developed and calibrated a new multi-scale computational modeling approach that incorporates the spatial patterning of key subcellular properties including cell contractility and cell division dynamics (Fig. 2B). This model for studying regulation of the subcellular features of morphogenesis of the growing *Drosophila* wing imaginal disc (Fig. 2D, E) also incorporates for the first time the dynamics of interkinetic nuclear migration within a pseudostratified epithelia (Fig. 3).

**Figure 3.**
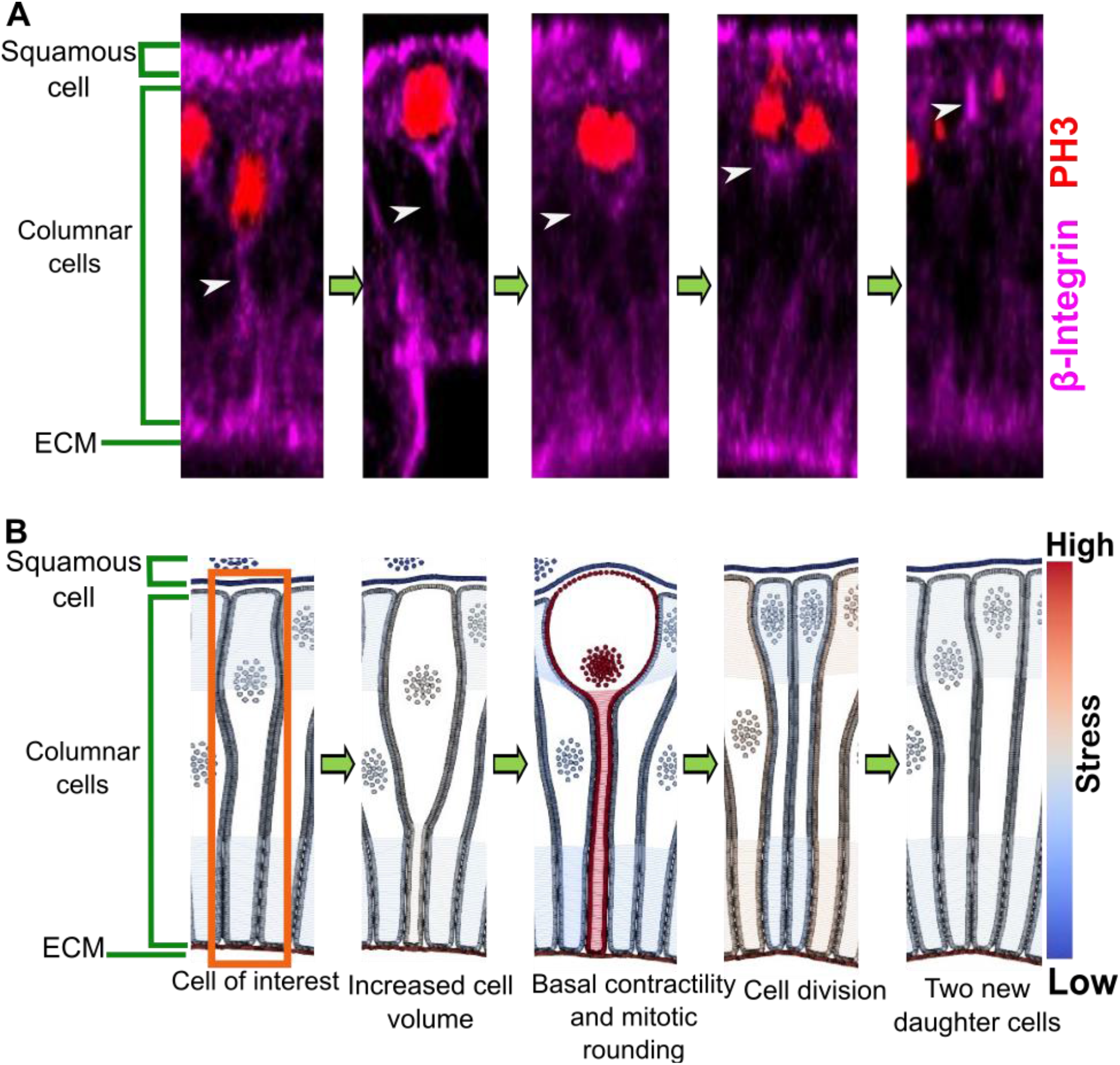
Subcellular element computational snapshots of a dividing cell capture the interkinetic nuclear migration and mitotic rounding process along with experimental representation. **(A)** A columnar cell in a wing disc cell demonstrating the phases of mitosis and interkinetic nuclear migration. White arrows are showing the Actin tails. **(B)** A simulated columnar cell undergoing division. The basal surface of the cell is connected to the ECM while the apical side adheres to a squamous cell.

Further, we report that key experimental observations of the dynamics of tissue shape changes across the final stages of larval wing disc development can be controlled by specific hypothesized mechanisms based on model simulations combined with experiments (Fig. 4, 5, 6). Analysis of experimental data reveals that an increase in local ratio of apical to basal levels of pMyoII leads to a decline in basal tissue curvature and an increase in thickness of the pouch cells specific to the medial domain (Fig. 4, 5). Experimentally we also observed diminishing of pMyoII at the basal-medial domain of the pouch with age of disc. Patterning of contractility levels within the model simulations similar to experimental actomyosin contractility profiles recapitulates the shape changes observed experimentally providing additional validation for the mechanism. Diminishing basal contractility within the model also pushed the nuclei in the medial domain towards the basal membrane (Fig. 6). This model prediction was validated through experiments where we observed an increase in basal migration of nuclei with age of the disc. Experimental confirmation of simulation predictions also led to discovery that triggering proliferation through the expression of the constitutively active form of the insulin receptor (InsR^CA^) in the posterior compartment of the wing disc increased compartment-specific basal epithelial curvature (Fig. 7). Lastly, we experimentally demonstrated that shape changes in Dpp receptor Thickveins (Tkv) mutants can be explained through a mechanism of dual regulation in apical and basal patterning of pMyoII and cell proliferation (Fig. 8). Overall, this work proposes and tests a novel mechanism of tissue growth control in a model pseudostratified epithelium that balances the competing effects of proliferation and cytoskeletal regulation to regulate tissue shape and size (Fig. 9).

**Figure 4.**
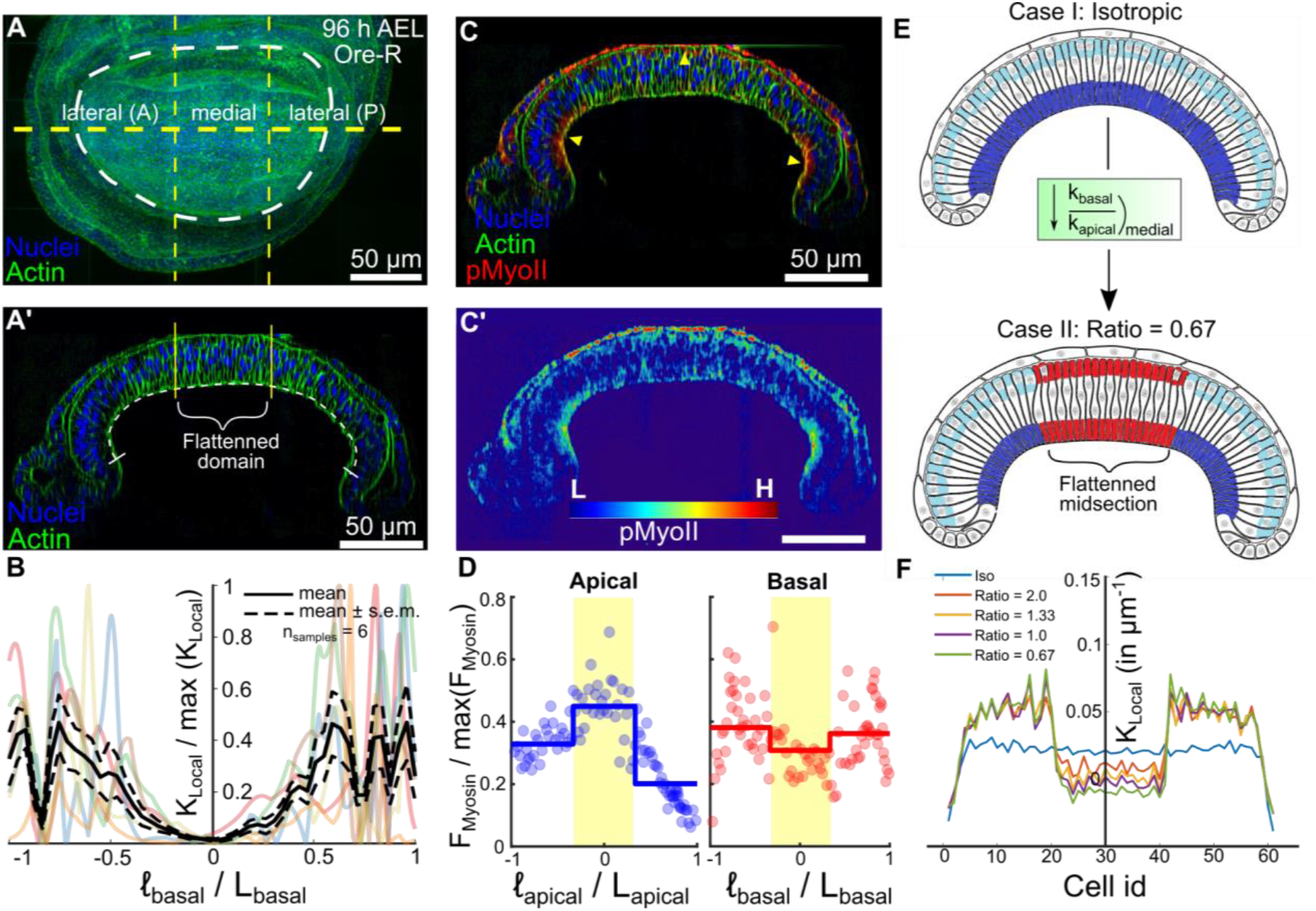
Spatial patterning of cell mechanical properties defines emergent tissue geometric features. **(A-A’)** Reference apical and cross-sectional views of a wild-type wing imaginal disc at early 3rd instar larval stage (96 h AEL). Nuclei are stained with DAPI and Actin with Phalloidin. **(B)** Plot showing the normalized curvature of the basal surface of the tissue calculated for six different samples. Normalization was carried out by dividing the local curvature (κ_local_) with the maximum value of the local curvature (max(κ_local_)). **(C)** Pattern of phosphorylated non-muscle Myosin II (pMyoII). (**C’)** A heat map of pMyoII distribution. Color-code represents lower (L) to higher (H) intensity of pMyoII.**(D)** Plots quantifying the fluorescence intensity of pMyoII along the basal and apical surfaces. The surfaces were further divided into 3 distinct subregions that included the two lateral and the middle portion of the cross-section. The solid lines are indicative of the region-wise averaged pMyoII intensity. **(E)** The color code indicates how apical and basal contractility parameters are specified. Different colors (red v.s. blue) represent the assignment of contractile spring coefficient based on the location of the model wing disc. The tints of color here represent the difference in the magnitude assigned such that lighter color indicates a lower contractile spring coefficient. Simulations demonstrate that isotropic patterning of the actomyosin contractility fails to generate the experimentally observed flat midsection. **(F)** Local basal curvature of the model tissue provided by different ratios of the basal-to-apical contractility. The ratio depicted here is for the midsection of the wing disc.

**Figure 5.**
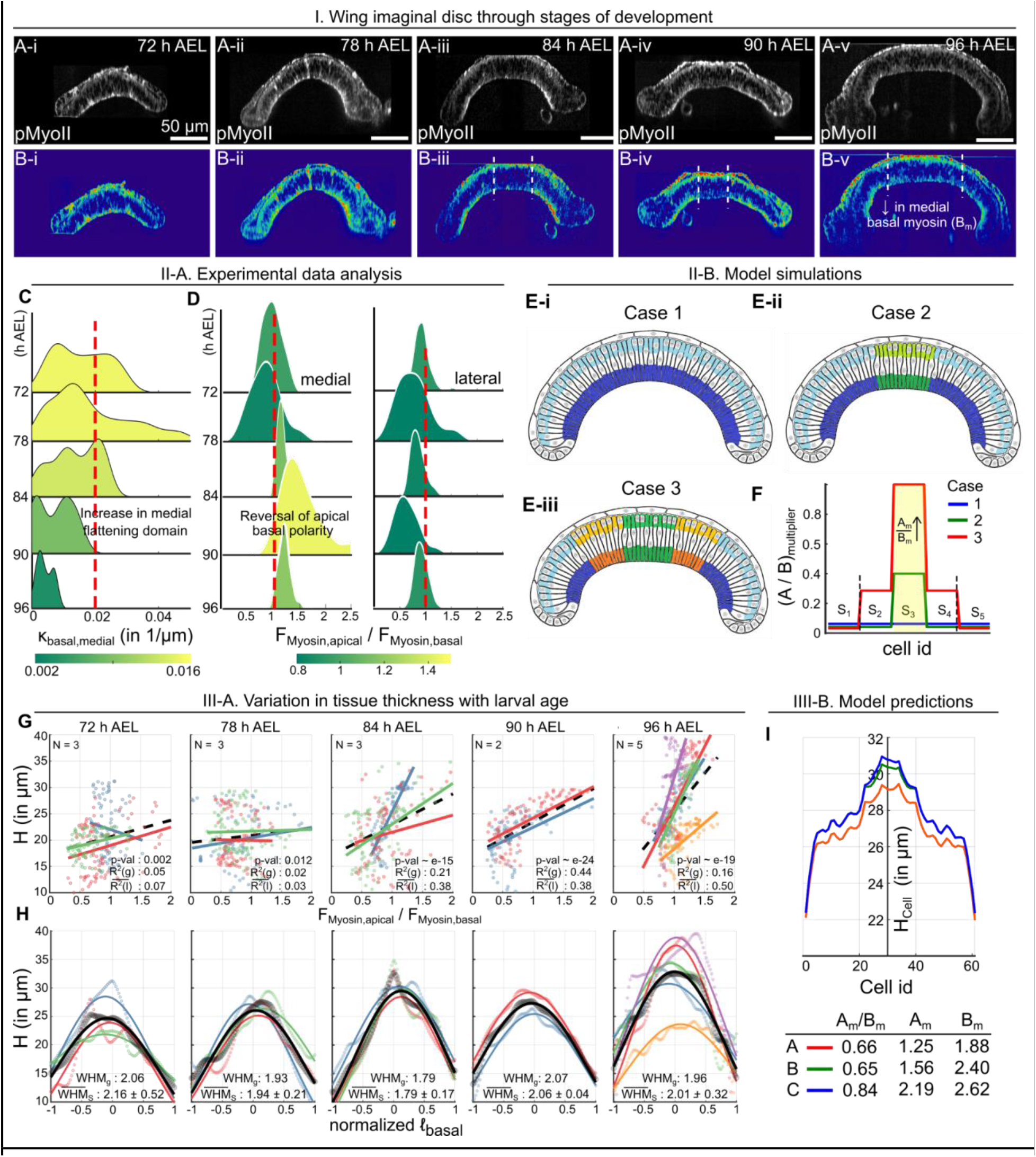
Local curvature of the basal epithelia anti-correlates with the apical to basal ratio of pMyoII in wing imaginal disc across multiple developmental stages. **(A i-v)** Fluorescence describing the localization of pMyoII within the wing imaginal disc at different developmental stages. **(B)** A heatmap distribution of pMyoII **(C)** Variation of the local curvature of the basal epithelia was represented as a kernel density estimate. Additionally, ridgeline plot was used to visualize the variation of curvature along the medial-basal surface for wing discs belonging to different developmental stages. **(D)** The DV optical section was discretized into 90 cells using the quantification pipeline described in Supplementary Fig. 12. Based on the location of these discretized cells along the apical and basal surfaces they were further subdivided into cells belonging to the medial and lateral domains of the pouch. Ratio of apical to basal levels of pMyoII were calculated for the cells. The corresponding distribution of these ratios of the medial and lateral subgroups were plotted as kernel densities. Ridgeline visualizations were next used to visualize the transition of apical to basal levels of myosin for wing discs of different ages. **(E)** Simulation results with different profiles of contractility in both apical and basal sides. The colors indicate the transition of contractility following simplified myosin dynamics. In terms of contractility strength, blue > orange > green, while the tints within the same color indicate the difference in the apical and basal contractility within each individual cell. The lighter color here indicates reduced contractility. **(F)** Transition of the apical and basal contractility used in the simulation. Following the pattern observed in the experiments, the activity of myosin gradually becomes more apical-bound in the midsection generating stronger contractility, while the reduced basal-bound myosin activity leads to the stagnation of basal contractility. **Local tissue height increasingly correlates with the ratio of apical to basal levels of pMyoII. (G)** Variation of tissue height with the ratio of apical to basal levels of pMyoII has been plotted. Different colors in the scatter represent different samples. A multilinear regression has been done to fit individual models to sample-specific data and is color-coded in a similar way to the raw data points. A solid line represents a linear model fit considering all the data from multiple samples. **(H)** Local tissue height evaluated using points on the basal surface were plotted against the normalized distances from the AP axis. A Gaussian equation was fit to the data and its WHM has been reported. **(I)** Height profiles in the tissue are plotted against the cell id (from left to right of tissue) for varying ratios of apical and basal contractility in the pouch medial domain.

**Figure 6.**
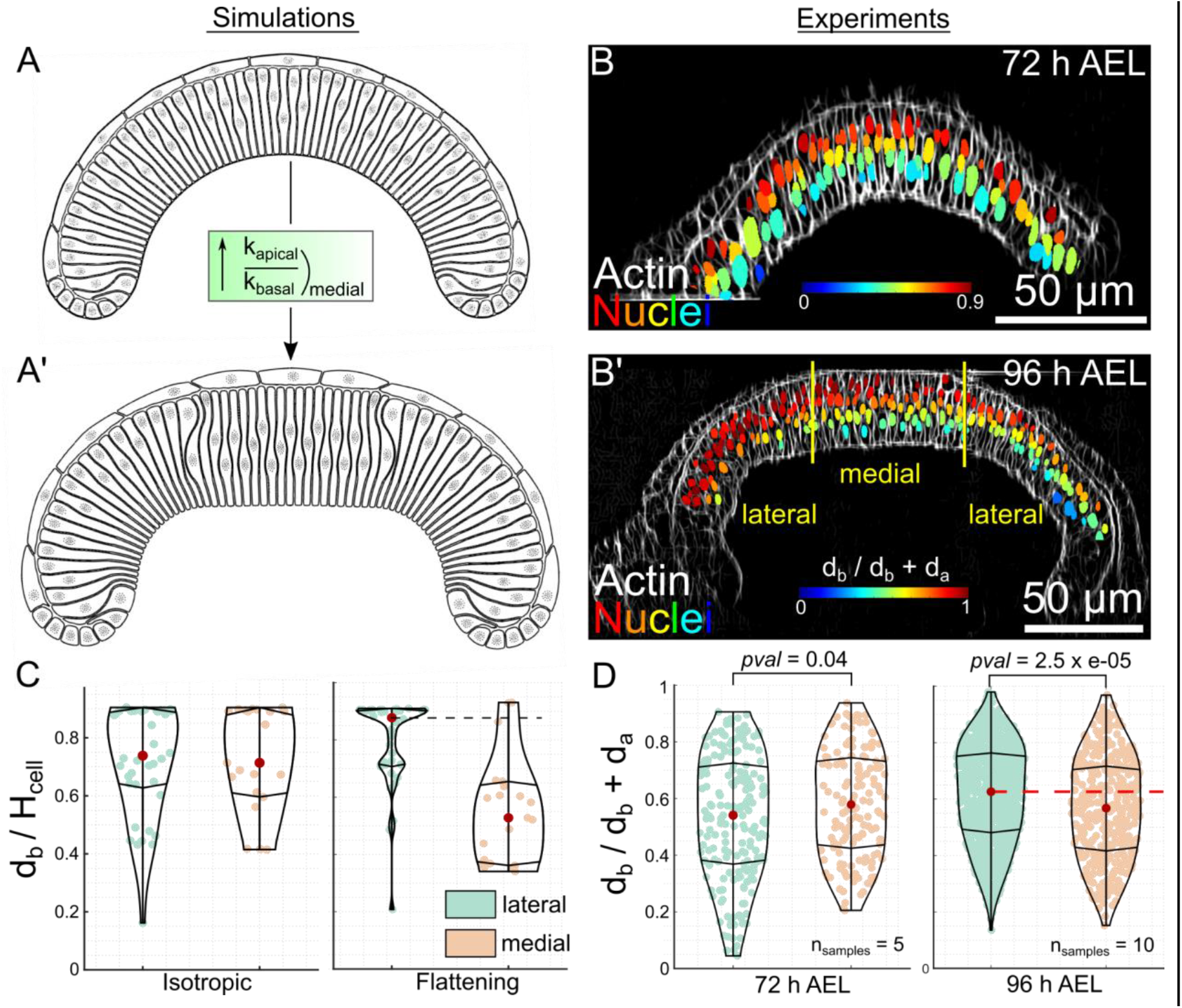
Higher levels of apico-central contractility results in a basal bias of the position of nuclei. **(A)** Nuclear positioning in a uniformly patterned actomyosin profile is distributed uniformly within each columnar cell. **(A’)** A non-uniform actomyosin patterning generates a flatter midsection which results in a shift of nuclear positions toward the basal side in the midsection. **(B-B’)** Optical sections along the DV axis for discs belonging to 72 and 96h AEL were used to mimic the patterning of contractility in the model simulations. Nuclei have been color-coded with respect to their distance from the basal surface. **(C)** Box-violin plot showing the differences in model-predicted relative nuclear positioning with respect to the basal surface in medial and lateral regions of the pouch. Relative nuclear positioning is defined as a ratio of the distance of nuclei from the basal surface and the corresponding cell height. **(D)** Box-violin plot showing the experimentally observed differences in relative nuclear positioning with respect to the basal surface in medial and lateral regions of the pouch. Relative nuclear positioning is defined as a ratio of the distance of nuclei from the basal surface and the corresponding cell height.

**Figure 7.**
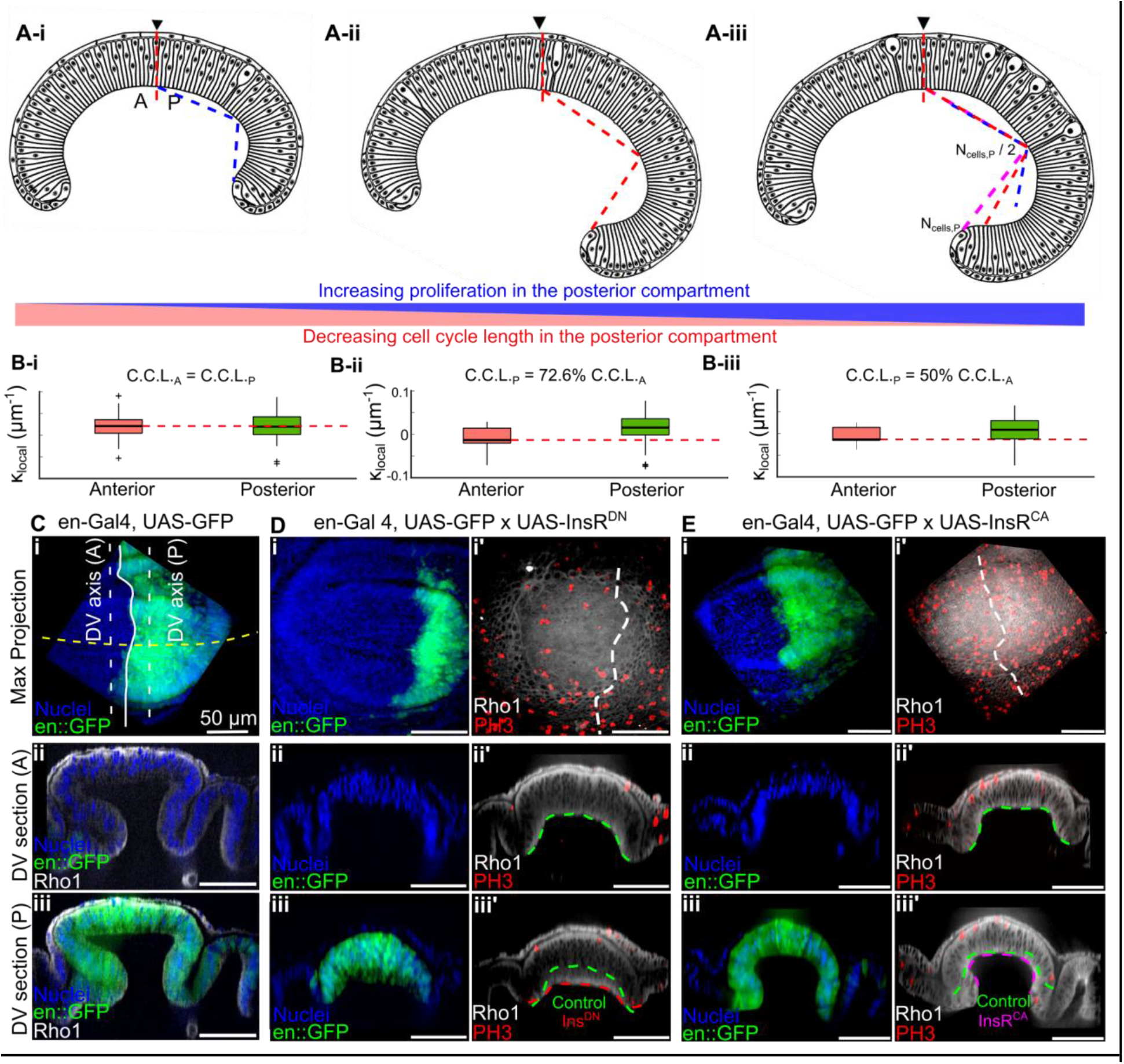
Stimulating cell proliferation by activated insulin signaling increases tissue curvature. Comparison of the local basal curvature between simulated wing disc cross sections with varying proliferation rates between the anterior (left) and posterior (right) compartments of the simulated tissues. These simulations assumed myosin activity is elevated on the basal side of the columnar cells. **(A, B-i)** Equal proliferation rates in both compartments. **(A, B-ii)** Cell cycle duration in the posterior compartment is set to be approximately 72.6% of the cell cycle in the anterior compartment. **(A, B-iii)** Posterior compartment of the wing disc has a cell cycle duration of approximately 50% of the cell cycle duration on the anterior compartment. **(B)** With increased cell proliferation, an overall increase in basal curvature is observed both qualitatively and quantitatively. The left box and error bar represent the data collected from the anterior side of the wing disc, and the right box and error bar represent the data **(C) Top-bottom**: Maximum intensity projection, and reslices parallel to the AP boundary in the anterior and posterior compartments for the control samples. **(D)** Similar data representation as C for the samples expressing dominant negative insulin receptor InsRDN in the posterior compartment. **(E)** Similar data representation as C for the samples expressing InsRCA in the posterior compartment. Fluorescent labels have been indicated on the bottom left corner and the 50 micron scale bar on the right.

**Figure 8.**
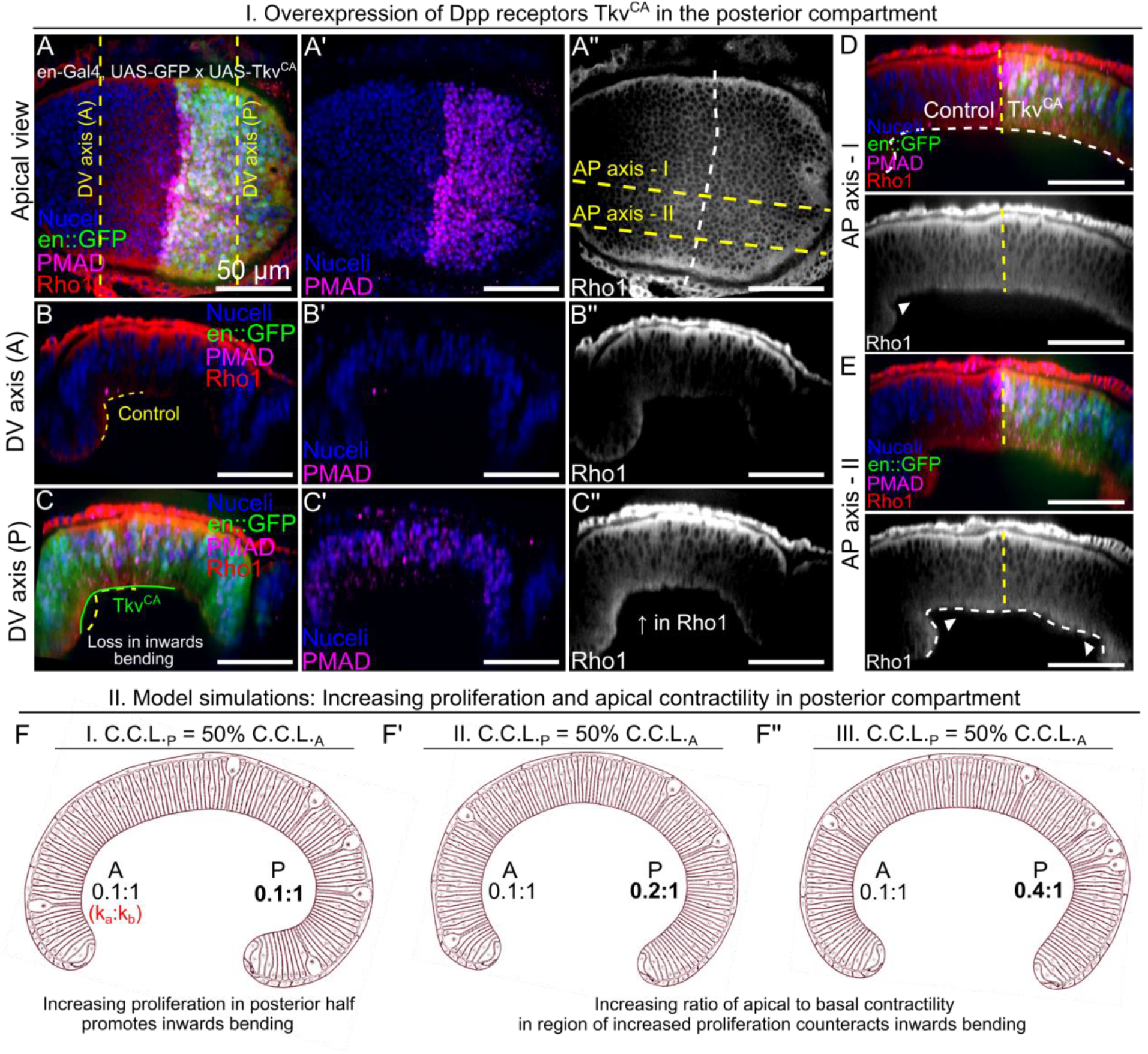
Dpp balances local curvature through joint regulation of proliferation and Rho-mediated actomyosin contractility. **(A-A’’)** Apical view of wing imaginal disc expressing a constitutively active form of Tkv receptors in the posterior compartment. Fluorescent labels are indicated in the bottom left corner and the scale bar is 50 microns. **(B-B’’)** Optical section along the anterior-posterior boundary taken in the anterior compartment of the wing disc. **(C-C’’)** Optical section along the AP boundary taken in the posterior compartment of the wing disc. (**D-E)** Optical sections along the AP axis taken at two separate locations represented by dashed yellow lines in A’’. **(F-F’’)** Comparison of the local basal curvature between simulated wing disc cross sections with varying ratio of apical-basal contractility between the anterior (left) and posterior (right) compartments of the simulated tissues. For these simulations, the posterior half of the wing disc proliferates twice as fast as the anterior half.

**Figure 9.**
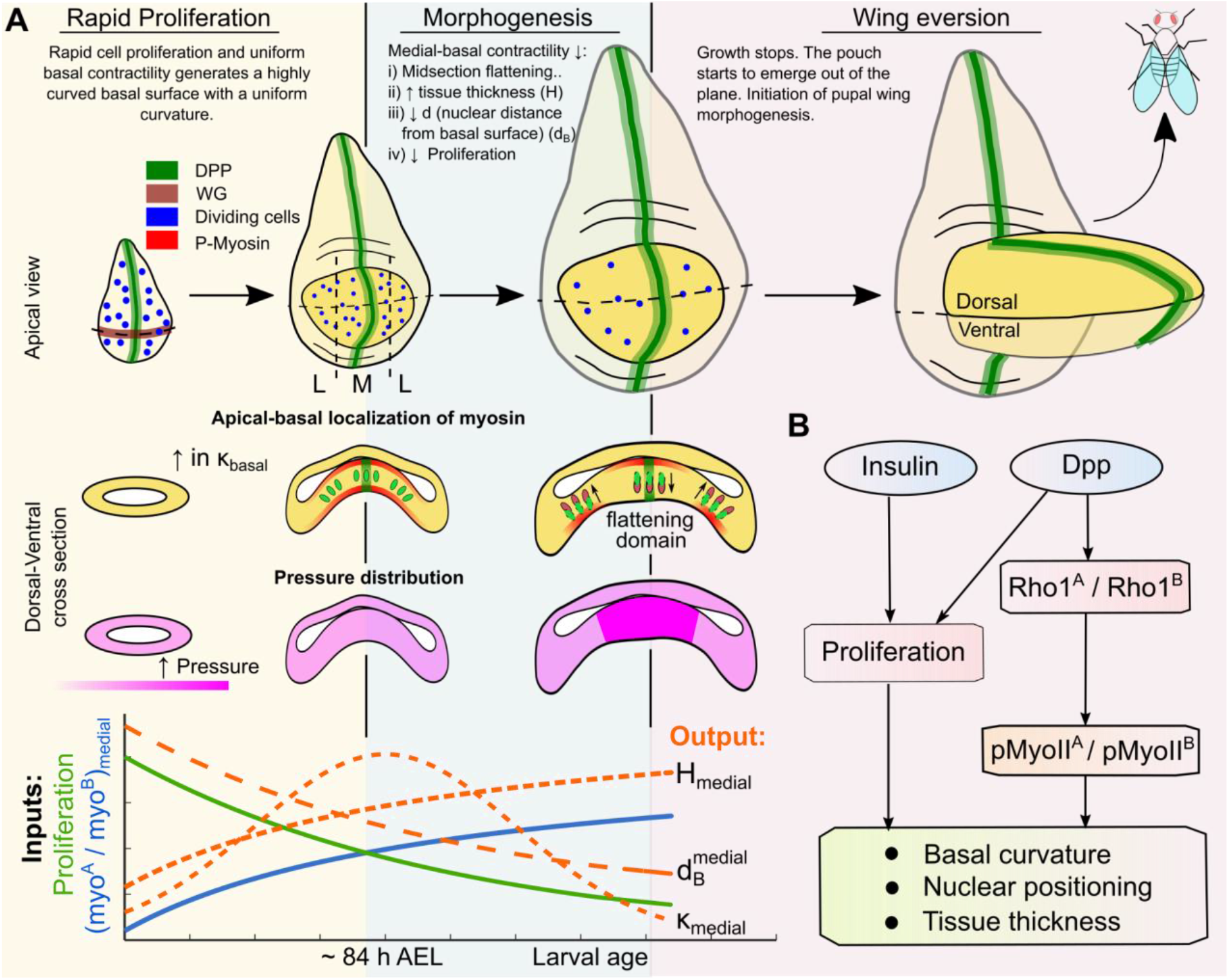
Integration of cell proliferation and cell mechanics regulation defines the shape of the growing wing disc. **(A)** Dynamics of tissue curvature directed by BMP/Dpp signaling. **(B)** Growth regulators come in two classes. As the first class, growth promoters such as insulin impact proliferation, which results in an enhancement of the natural curvature of the wing disc. In the second class, growth promoters such as the morphogen Dpp balance the competing impacts of cell proliferation and cytoskeletal regulation to reduce the net curvature of the tissue while still promoting growth.

## Results

### Wing disc growth termination includes both cell height thickening and tissue flattening

To understand the interrelationships between tissue growth and morphology, we quantified wing imaginal disc shapes taken from larvae over multiple time points leading up to pupal stages (72-96 h after egg laying, AEL). Analysis of basal curvature of the optical sections taken along the DV boundary revealed a flattening in the center of the wing disc (medial domain). This flattened section exhibiting low local curvature increases in width (Fig. 1B). The local thickness profiles of the cross sections show an increase in cell height (Fig. 1C). Along with the bulk shape changes, we also report a basal shift of nuclei in the pouch midsection as the disc grows in size (Fig. 1D). Our results indicate an apical shift of phosphorylated non-muscle myosin II (pMyoII) as the discs grow (Fig. 1E). With these cytoskeletal dynamics, cell proliferation decreases in agreement with the previous studies^15,31,32^ (Fig. 1F), (Supplementary Fig. 1). These observations of dynamic changes in cell-level processes such as contractility and proliferation suggest possible coordinated regulation between cell growth and proliferation and the cell mechanics that define the emergent shape of the tissue.

### Predictive multiscalar subcellular element model simulations of proliferation and cytoskeletal regulation in a pseudostratified epithelium

To develop a predictive model that explains the shape changes in the wing imaginal disc as it grows in size, we developed a new subcellular element model focusing on the A-P cross-section (Fig. 2A-ii). In principle, this approach can also capture morphology along the D-V cross-section (Fig. 2A-i) and is a focus of future efforts. In our previous work^33^, it was assumed that actomyosin contractility is patterned uniformly across the apical and basal surfaces of the pouch. However, our experimental analysis (Fig. 2B), (Supplementary Fig. 2) of major cytoskeletal regulators reveals anisotropy in the spatial patterning of mechanical regulators such as Actin, phosphorylated non-muscle myosin II (pMyoII), and Integrin across the AP axis. For instance, Actin, pMyoII, and Integrin are more concentrated at the lateral basal ends than in the medial domain of the pouch at later stages of development (84 h AEL and later) (Fig. 2B). To test the significance of this asymmetry, we developed new simulations that include spatial variation of cell-level mechanical properties.

A second key innovation in this study is that this is the first subcellular element model that provides a detailed simulation of cell proliferation and growth throughout the stages of development. Owing to the high computational cost of subcellular element model simulations, previous efforts only allowed for investigations into shape regulation at a single developmental stage and without spatial patterning of individual cell properties. Here, we created a platform that simulates cell divisions and cell growth throughout organ development. To model cell division, we simulated the multiple stages of interkinetic nuclear migration (Fig. 2C, D, Fig. 3). During division the cell experiences narrowing of the basal section due to actomyosin contractility and depolymerization of the apical contractile springs. During this time, the nuclei migrates towards the apical surface of the pouch. After division, the apical and basal contractile springs of the two new daughter cells are restored to pre-division values (Fig. 3B). Additional implementation details are provided in the methods section of this paper. To ensure tissue integrity, we postulate that the ECM gets pulled up as the nucleus migrates towards the apical surface during cell division. Interestingly, we discovered that this occurs as shown by the presence of both Integrin and Collagen IV, key components of ECM-cell adhesion, along the actin tail of dividing cells (Fig. 3), (Supplementary Fig 3). Finally, we developed a pipeline to calibrate the subcellular element model with the quantified experimental data to provide the agreement of local shape features such as nuclear positioning, basal curvature, and tissue thickness (Fig. 2E).

### Increasing the ratio of apical to basal contractility flattens the center of the wing disc

As noted above, the basal curvature is higher at the lateral ends as compared to the medial domain of the pouch (Fig. 4A’, B). We hypothesized that anisotropy in the patterning of cytoskeletal regulators at a cellular level accounts for such tissue shape changes. To test this, we employed the quantification pipeline for fixed and stained samples for pMyoII (Fig. 4C-C’), Integrin, and Collagen IV, one of the key components of ECM (Supplementary Fig. 2). Interestingly, we observed a high expression of Integrin and Collagen IV at the lateral ends of the basal side of the epithelium as compared to the midsection (Supplementary Fig. 2). This likely represents an increase in adhesion between the cells and ECM at increasing distances from the center of the pouch. We also reported an anisotropic patterning of pMyoII across both the apical and basal surfaces (Fig. 4D). Myosin was more localized at the apico-medial and lateral ends of the basal surface. This observed anisotropy in the patterning of cytoskeletal regulators at both the apical and basal surfaces suggests that achieving a quantitative agreement between experimental data and computational simulations must incorporate spatial variation in the model parameters that define the forces acting on the cell at the subcellular level.

We take as a starting point our previous, calibrated computational subcellular element model that qualitatively explains the curved wing disc shape and which only utilizes spatially homogeneous mechanical properties^33^. However, this approach cannot explain the non-uniform bending profile that develops later in development with flattening in the center of the pouch. Here we find that flattening in the center of the pouch can be explained by regulating the spatially patterned balance of apical and basal contractility. Tissue shapes were simulated by varying values of parameters describing cell mechanical properties in the model including the parameters responsible for the apical and basal actomyosin contractility (*k*_*api,cont*_ and *k*_*bas,cont*_), the Integrin-based adhesion between the cells and the basal ECM (*k*_*adhB*_) and the basal ECM stiffness (*k*_*ecmc*_).

All simulations started with the same flat tissue shape. By assuming homogeneous contractility on both apical and basal sides of individual cells with higher strength at the basal side, the tissue evolved into a bent shape with a globally uniform curvature (Fig. 4E, Case I).

With lower contractility in the center on both apical and basal sides, the tissue shape was flatter in the middle (Fig. 4E, Case II), which is more similar to the shape observed in experiments. We also showed that a patterned cell-to-ECM adhesion and ECM stiffness did not have a significant impact on the basal curvature in simulations (Supplementary Fig. 4). Our experimental data together with the simulation results suggest that nonhomogeneous actomyosin contractility is sufficient to achieve the tissue shape with non-uniform curvature where the medial section is flat. However, at later stages, the pMyoII_apical_/pMyoII_basal_ ratio exceeds one for the medial domain indicating a reversal in polarity of apical-basal pMyoII in the central region of the pouch. The increase in the ratio of apical to basal levels of myosin in the medial domains qualitatively correlates with a decrease in local basal curvature (Fig. 5C, D).

### Local curvature of the basal epithelia anti-correlates with the apical to basal ratio of pMyoII across multiple developmental stages

To further test the hypothesis that higher apical contractility flattens the midsection, we extended the experimental analysis of patterned myosin and basal curvature of wing discs from multiple developmental stages (Fig. 1B, Fig. 5A, B). At earlier stages of development, the curvature of the basal surface is uniformly distributed across the anterior-posterior axis. However, with an increase in both age and size of the disc, the midsection starts to flatten out with respect to the more bent anterior and posterior ends (Fig. 1B, Fig. 5C). Moreover, the spatial width of the flattened region also increases with age. Next, we quantified the ratio of apical to basal levels of pMyoII in the medial and lateral domains of the pouch. The distribution of this ratio is represented as a ridge map of kernel density estimates (Fig. 5D). At earlier stages of development, pMyoII is more basal in both medial and lateral domains as the ratio of apical to basal levels of pMyoII is predominantly less than one (Fig. 5A-i-ii, B-i-ii). However, at late stages of morphogenesis (> 78 h AEL), pMyoII in the medial domain of pouch is mostly apical (Fig. 5A-iii-v, B-iii-v).

The computational model simulations were run to test the hypothesis that this change in the ratio of apical to basal pMyoII concentrations was sufficient to capture the dynamics in shape changes as the wing disc grows in size. With homogeneous contractility at both apical and basal sides of individual cells, the tissue developed into a bent shape with uniform curvation (Fig. 5E-i, Supplementary Video 1). By lowering the contractility strength in the tissue center at both apical and basal sides, the tissue was flattened in the middle (Fig. 5E-ii). The domain of flattened tissue expanded when further lowering the contractility strength at the center and introducing three different levels of contractility strength at both apical and basal sides following an increasing trend from the center to the boundary (Fig. 5E-iii). Therefore, the domain of flattened tissue depended on the region with low contractility strength marked by diminished basal-medial pMyoII levels.

### Strength of correlation between local tissue height and ratio of apical to basal levels of pMyoII increases with wing disc age

Average tissue height increases as the discs age from 72 h to 96 h of larval development (Fig. 5G, H). Since apical localization of myosin increases with age (Fig. 5D), we examined the extent it correlates with the increase in cell height observed in experiments.

To test if the ratios of apical to basal contractility are sufficient to regulate cell height, the model parameters *k*_*api,cont*_ and *k*_*bas,cont*_, specifying the apical and basal actomyosin contractility respectively, were varied systematically in computational simulations (Fig. 5I), (Supplementary Fig. 5). In the first set of simulations, we increased both *k*_*api,cont*_ and *k*_*bas,cont*_ in the medial domain of the pouch keeping the ratio of two constant. An increase in the contractility values in both the apical and basal surfaces of the pouch led to an increase in average tissue thickness. This increase in the tissue thickness is limited to only the medial pouch domain where the increase in the apical and basal contractility is introduced. In the next set of simulations, we increased the parameter values that specify apical and basal levels of contractility so that the ratio of the two is also increased. Similar to the previous case, the medial cell height increased (Fig. 5I).

We also studied the role of cell volumes in generating this gradient in cell height. Cell volumes in the medial pouch domain were increased as compared to the cells in the lateral ends. This increase in cell volume is mediated by linearly increasing the value of the equilibrium cell volume 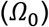 at each simulation time step (see Methods). A thickening in the medial pouch domain was observed upon increasing the cell volume in the central pouch region (Supplementary Fig. 6). However, previous studies do not report patterning of cell volumes along the DV axis of the pouch. Our variation of parameter values suggest that local tissue height is primarily regulated through subcellular localization of actomyosin contractility.

Since we already established that the ratio of apical to basal levels of pMyoII regulates cell height, we next studied the correlation between this ratio and local tissue thickness (Fig. 5G). For discs belonging to early stages of development (< 84 h AEL), the ratio of apical to basal pMyoII levels poorly correlates with the average tissue height (R^2^ values <0.05).

Interestingly, the quality of fit and R^2^ values increase at later development stages suggesting that the correlation between apical myosin concentration and tissue height increases as the organ matures. At 96 h AEL, the averaged R^2^ value increases to 0.50 (n=5 samples) (Fig. 5G). Of note, while cell height does not correlate with the ratio of apical to basal levels of pMyoII or younger discs, a gradient in tissue height across the DV pouch section was still observed (Fig. 5H). This analysis of the experimental data reveals that although the local tissue thickness correlates strongly with the ratio of apical to basal pMyoII for discs only belonging to later stages of development, it is not the single unique factor that generates and sustains this patterning of cell height.

### High levels of apico-central contractility push the nuclei basally

Additionally, the apical and basal actomyosin contractility determines positioning of the nuclei in cells in model simulations of the wing disc. When the basal contractility is higher compared to the apical contractility, the positioning of the nuclei shifts toward the apical end of each cell (Fig. 6).

In the case of a uniform patterning of the actomyosin profile, the nuclear positioning (or apico-basal distribution) shows marginal difference across the whole wing disc (Fig. 6A, C). To generate the distribution of the overall nuclear positioning closer to the basal end of each cell in the midsection, as observed in experiments, the apical contractility needs to be comparable to the basal contractility (Fig. 6A’, C). We analyzed the positioning of nuclei in wing imaginal discs from 72 hours and 96 h AEL to validate the model prediction that nuclei are shifted basally in the center of the wing disc as development progresses. As discussed in the previous sections (Fig. 5), an early-stage wing imaginal disc mimics the case where the patterning of contractility parameters within the model is isotropic. On the other hand, wing discs from a later developmental stage have been used to study the case of patterned contractility. The discs were fixed and stained with dyes to fluorescently label Actin and Nuclei (Fig. 6B-B’).

It can be clearly seen that at the initial stages, the distribution of nuclear positioning seems uniform across the DV cross-section (Fig. 6B, D). However, with age, the nuclei in the medial domain are more basally located compared to those in their lateral counterparts (Fig. 6B’, D). This observed shift is in qualitative accordance with the model predictions. In summary, our results in this section indicate a strong correlation between the apical-basal polarization of myosin and nuclear positioning within the epithelium.

### Stimulating cell proliferation through increased insulin signaling enhances local basal curvature

Computational model simulations predict that increased cell proliferation without concurrently changing the uniform patterning of the actomyosin contractility leads to an increase in the basal curvature (Fig. 7). The initial geometry of the wing disc specified in simulations reflects the curvature observed in experiment wing discs at approximately 72 h AEL. To test how cell proliferation impacts tissue geometry, we increased the proliferation rate in the posterior side of the model wing disc more than the proliferation rate on the anterior side. In particular, we set the posterior-to-anterior cell cycle length ratio of 72.6% for Fig. 7A-ii and 50% for Fig. 7A-iii in contrast to a spatially homogeneous cell division rate for the “wild type” control simulation (Fig. 7A-i). With increased cell proliferation driven by a reduced cell cycle length, the simulated tissue developed an overall increase in the local basal curvature (Fig. 7B).

Qualitatively, the posterior side of the simulated wing disc also shows a more “inward” curving (positive basal curvature) as the rate of proliferation increases (Supplementary Video 2). To test the model simulation’s predictions that proliferation increases curvature, we expressed dominant-negative (DN) and constitutively-active (CA) forms of *Drosophila* Insulin receptors (InsR) in the posterior compartment of the wing disc using the engrailed-Gal4 drive (Fig. 7C-E). Downregulation in Insulin signaling activity through the expression of InsR^DN^ led to a reduction in the number of mitotic cells on the posterior side of the compartment, as expected^34^ (Fig. 7Di-i’). InsR is a regulator of proliferation in epithelial cells as shown by the increase in cell proliferation with overexpression of InsR^CA^ (Fig. 7Ei-i’).

We tested how this proliferation impacts tissue geometry by analyzing tissue cross-sections parallel to the AP axis in the control and perturbed tissue sections. For the control engrailed driver, the AP optical sections are identical indicating a symmetry in the patterning of geometrical properties along the DV axis (Fig. 7C-(ii, iii)). Upon expression of Ins^DN^, the AP reslice taken in the region of perturbation i.e. posterior side becomes flatter compared to its anterior counter-half. The basal surface of the control domain is highlighted with a green dashed line and overlaid with the red dashed line representing the basal surface of the AP cross-section expressing InsR^DN^ (Fig. 7D-(ii’, iii’)). Similarly, overexpression of InsR^CA^ resulted in an increase in basal curvature (Fig. 7E-(ii’, iii’)). We also report no changes in the patterning of Rho1, an upstream regulator of pMyoII upon expression of both InsR^DN^ and InsR^CA^. This suggests that proliferation stimulated by insulin signaling promotes curvature in an actomyosin contractility-independent manner.

### Dpp maintains basal curvature through a balance of growth and Rho-mediated actomyosin contractility

Next examined the roles of morphogens in regulating cell mechanics during tissue growth. Dpp regulates cell growth, cell fate specification and patterning, and Rho GTPase activity within the wing imaginal disc. Regulation of RhoGTPase by Dpp patterns actomyosin contractility and impacts cell height changes in the pouch cells^7^. However, how Dpp signaling jointly regulates the balancing of local basal curvature, and cell height remains unclear. To investigate this, we perturbed Dpp signaling pathway activity in the posterior compartment. This enables comparison of tissue architecture with an internal wild-type anterior compartment as the control. As expected, expression of constitutively receptor Thickveins (Tkv^CA^) markedly increased phosphorylation of the transducer MAD (PMAD) (Fig. 8A’).

Constitutive active Dpp signaling stimulates overgrowth phenotype in the pouch^35,36^, and hence an increase in local basal curvature upon expression of Tkv^CA^ was expected. However, we report a loss of basal curvature and inwards bending near the lateral ends of the pouch (Fig. 8B, C, D, E). To study why this Dpp triggered proliferation results in loss of inwards bending, we first computationally increased proliferation in one half of the pouch by 50% and then varied the apical and basal contractility parameters (*k*_*api,cont*_ and *k*_*bas,cont*_) in the region of perturbation (Fig. 8F-F’’). The rationale behind such a computation is based on the idea that Dpp regulates pMyoII through Rho GTPases. The increasing ratio of apical to basal levels of contractility resulted in a loss of inward bending at the lateral end of the pouch having more proliferation. As an experimental validation results from our Rho1 antibody immunohistochemistry data also show an increase in Rho1 fluorescence in the posterior compartment (Fig. 8B’’, C’’). Expression of Tkv^CA^ is also known to increase pMyoII expression in wing imaginal discs^7^. Taken together our study suggests that increased apical contractility upon expression of Tkv^CA^ counteracts the proliferation mediated inwards bending of the wing imaginal disc.

Analysis of the experimental data also suggests an increase in cell height upon expression of Tkv^CA^ in the wing imaginal disc^7^ (Supplementary Fig. 7). This agrees with simulations that show an increase in columnar cell height upon an increase in both apical and basal contractility (Fig. 5I).

## Discussion

One of the most important unresolved problems in developmental biology is how a tissue’s final shape and size are coordinated through regulation of both cell proliferation and the cellular cytoskeleton. The relative contributions of cell proliferation and cell mechanics to the final morphology of an organ is context dependent. In some situations, proliferation and morphogenesis are separated into nonoverlapping temporal stages. For example, cell proliferation halts before gastrulation begins in the fly^37^. This strategy is likely important when developmental speed is of the essence and the tissue cannot reach a pseudo-steady state before a new developmental event occurs. In other contexts, proliferation and cell shape changes occur simultaneously as is the case during the growth and morphogenesis of the vertebrate optic cup^38^.

We introduced in this paper a new multi-scale computational model to quantitatively investigate the emergent features of morphogenesis through close integration with experiments (Fig. 1, Fig. 9A). Namely, we developed and calibrated a subcellular element (SCE) model of both tissue growth and morphogenesis that incorporates the spatial patterning of key subcellular properties and cell division dynamics (Fig. 2). We show how key characteristics of global tissue architecture such as the local curvature of the basal wing disc epithelium, cell height, and nuclear positioning are jointly regulated through spatiotemporal dynamics in the localization of phosphorylated non-muscle Myosin II (pMyoII), which in turn is downstream of key growth and developmental signaling pathways such as Dpp/BMP^39^ (Fig. 1, 4, 5). A uniform and high basal contractility during earlier developmental stages generates a highly curved tissue with homogeneous basal curvature and positioning of nuclei. As the disc grows in size, the relative concentration of pMyoII concentration shifts toward the apical side in the pouch medial domain. The change in polarization of pMyoII concentration is reflected through the flattening of the midsection along with an increase in pouch thickness and basal migration of nuclei (Fig. 6). Combining these experimental observations with the detailed simulation results yielded a conclusion that the apically activated pMyoII, followed by the reduction in basal contractile forces, increases the flatness in the medial region of the pouch (Fig. 9).

An analysis of growth dynamics within the tissue both in experiments and simulations revealed the importance of proliferation in regulating the local curvature of the basal epithelium. An increase in cell proliferation both computationally and experimentally through the expression of constitutively active Insulin receptors (InsR^CA^) increased the median basal curvature and inward bending of lateral domains (Fig. 7). Finally, we show the dual role of Dpp, a key morphogen, in regulating epithelial morphogenesis through the balancing of two separable mechanisms: a) cell proliferation, and b) spatiotemporal dynamics of apical-basal polarization in myosin (Fig. 8). This proposed model is able to explain how shape changes are a reflection of both proliferation and actomyosin contractility patterning in the wing imaginal disc during larval development. These shape changes likely are needed to prepare the wing imaginal disc for later pre-pupal and pupal stages of development^40^ (Fig. 9).

These findings reflect similar observations in other developmental contexts. The dynamic and autonomous morphogenetic process of formation of the optic cup, i.e., retinal primordium, is reminiscent of the tissue flattening that occurs before subsequent eversion of the wing imaginal disc^38^. During the formation of the mammalian optic cup, the invagination happens in four stages. Flattening of the distal region of the initially hemispherical vesicle is observed in the first two stages. The angle at the hinge then begins to become narrower following which finally, in phase four, the neural retina epithelium started to expand as an apically convex structure, forming a cup via progressive invagination. This morphogenetic process, specifically the last two phases, resembles the tissue bending and flattening in the wing disc. Balancing mutually antagonistic morphogenetic processes is seen in mammalian development as well, including the developing mouse lens^41^.

A myriad of structural proteins act as an anchor to support the maintenance of nuclear positioning within a cell, and mutations impacting nuclear positioning can lead to a number of human diseases^42^. Previous studies identified cell density and local tissue curvature as factors that influence interkinetic nuclear migration during mitosis^8^. For instance, a flatter zebrafish hindbrain reduces apical migration during mitosis as opposed to a more curved retina^43^. However, the converse question of how cell division impacts overall tissue morphology is still incompletely understood. Here, we investigated how proliferation impacts curvature. We found that insulin-stimulated growth and proliferation accentuates the natural basal curvature of the wing disc epithelium (Fig. 7). These results raise new questions regarding the feedback loops between cell cycle control determined by gene regulatory networks and tissue morphology defined by “cell regulatory networks”^3^. How nuclear positioning during interphase impacts the cell cycle dynamics in pseudostratified epithelia is also unclear.

This work highlights how the relative balance between apical and basal contractility regulates the positioning of nuclei within wing imaginal discs (Fig. 6, Fig. 9A). Both simulations and experimental analysis of temporally staged wing discs reveal that declining basal contractility within the medial pouch domain causes nuclei to shift toward the basal side of the epithelium. Concurrently, a decline in cell proliferation occurs as the tissue matures. These two observations suggest a potential causal link between the effective migratory distance for nuclei and cell cycle duration. A reduced basal level of pMyoII may be associated with a reduced tendency to generate the contractile force needed to facilitate interkinetic nuclear migration.

As the disc grows in size, there is a change in patterning of key subcellular features such as actomyosin contractility (Fig. 5). However, the current model does not include an explicit coupling between the maturation state of the tissue and the regulation of contractility. For simulations of the tissue at different developmental stages, contractility parameters within the model have to be calibrated based on the experimental data. Morphogens such as Dpp and RhoGTPases^44,45^ regulate actin filament dynamics and myosin motor activity. Future investigations are needed to develop a coupled, mechanistic chemical signaling model that mechanistically couples cell differentiation state with tissue mechanics.

In this work, we assumed that the dominant contractile force occurs through contractile rings that grow laterally on the basal side of the cell during interkinetic nuclear migration. During cell division, the basal section of the dividing cell is still in contact with the basal ECM (Fig. 3). High localization of both Integrin and Collagen IV, a key ECM component, was observed in the actin tail and the cortical region of the mitotic nuclei (Fig. 3, Supplementary Fig. 3). However, this does not resolve whether the length of basal lateral edges are regulated during cell division and if the basal ECM is “pulled up” during interkinetic migration. These hypotheses remain to be confirmed experimentally. However, presence of Integrin and Collagen IV around the dividing cell does provide a preliminary evidence for ECM being “pulled up” during mitosis. Future developments of this model will require more detailed description of the interaction between the actin filaments and myosin motors such that the directionality of the contractile forces can be explicitly incorporated, and the mitotic rounding process along with cell division represented in full mechanistic detail. Overall, this study highlights how the dynamic balance of growth and shape control is differentially regulated by both morphogens and growth control pathways, leading to a general strategy governing organ morphogenesis that can tune the coupling between growth and tissue curvature (Fig. 9B).

## Methods

### Experimental and image analysis methods

#### Fly stocks and culture

*Drosophila* were raised in a 25 °C incubator with a twelve-hour light cycle unless specified otherwise. Virgins from Gal4 lines were collected twice a day from the bottles. Virgins were crossed with males that carry the indicated UAS line constructs in a ratio of female to male of 15:5. The crosses were staged for 4 hours to collect the correct aged larvae. The wild-type Oregon-R fly line is a long-standing stock in the Zartman lab acquired from the Niryakobi lab. The following transgenic stocks were obtained from Bloomington Drosophila Stock Center (BDSC): UAS-RyR^RNAi^ (BDSC#31540), UAS-InsR^DN^(BDSC#8253), UAS-InsR^CA^(BDSC#8263), UAS-Tkv^RNAi^(BDSC#31041), UAS-Tkv^CA^(BDSC#36537), Nubbin-Gal4 (BDSC#25754), Engrailed-Gal4 (BDSC#25752).

#### Immunohistochemistry

Wing imaginal discs were dissected in phosphate-buffered saline (PBS) from 72 h, 96 h, or 120 h AEL larvae in intervals of 20 min. Fixation was performed on dissected wing discs in ice-cold 4% paraformaldehyde solution (PFA) in PBS for 1 h in PCR tubes. Fresh PBT (PBS with 0.03% v/v Triton X-100) was used to rinse the wing discs three times immediately following fixation. PCR tubes containing wing discs in PBT were then placed on a nutator for 10 minutes at room temperature and then rinsed again with fresh PBT; this step was repeated for three nutation/rinsing intervals. PBT from after the third rinse was removed and 250 μL of 5% normal goat serum (NGS) in PBS was added to the PCR tube. Tubes were then agitated on a nutator for 45 minutes at room temperature. NGS was replaced with 250 μL of a primary antibody mixture prepared in a 5% NGS solution. Next, tubes were agitated on a nutator at 4 °C overnight. The following primary antibodies were used: Phospho-Smad1/5 (Ser463/465) (41D10) (1:300, Rabbit, Cell Signaling Technology #9516S), P-Histone H3 (1:500, Rabbit, Cell Signaling Technology #9701S), Phospho-Myosin Light Chain 2 (Ser19) (1:50, Rabbit, Cell Signaling Technology #3671S), α-Rho1 (1:10, Mouse, Developmental Studies Hybridoma Bank p1D9), Integrin betaPS (myospheroid) (1:5, Mouse, Developmental Studies Hybridoma Bank CF.6G11), α-Collagen IV antibody (1:5, Rabbit, Abcam ab6586). Three PBT rinses were performed the next day as was done after fixation. Tubes were then placed on a nutator for 20 minutes with fresh PBT and this was repeated for three nutation/rinsing intervals. After removing the PBT from the final rinse, 250 μL of a secondary antibody mixture corresponding to the primary antibodies along with DAPI, prepared in a 5% NGS solution, was added. Tubes were then agitated on a nutator for 2 hours at room temperature. The following dyes and secondary antibodies were α-Mouse Alexa Fluor™ 568 (1:500, Goat, Thermo Fisher Scientific A-11031), α-Rabbit Alexa Fluor™ 647(1:500, Goat, Thermo Fisher Scientific A32733), DAPI (1:500, Sigma Aldrich D9542), Fluorescein Phalloidin(1:500, Thermo Fisher Scientific F432).Three subsequent quick PBT rinses followed by three 20 minutes of agitation on the nutator with fresh PBT at room temperature were performed. Wing discs in PBT were left in the tube at 4 °C for an overnight wash. Four sets of double-layered scotch tape strips were used as spacers to create a square well. The spacers were positioned on the surface of a glass slide which prevents the coverslip from pressing on the wing discs. The wing discs were mounted within the well using Vectashield mounting medium and a cover slip was placed atop, aligned with the spacers.

#### Confocal Microscopy

The wing imaginal discs were imaged using a Nikon Eclipse Ti confocal microscope with a Yokogawa spinning disc and MicroPoint laser ablation system, and a Nikon A1R-MP laser scanning confocal microscope. For the two confocal microscopes, image data were collected on an IXonEM+colled CCD camera (Andor Technology, South Windsor, CT) using MetaMorph v7.7.9 software (Molecular Devices, Sunnyvale, CA) and NIS-Elements software, respectively. Discs were imaged throughout the entire depth of z-planes with a step size of 0.8-1 μm, depending on sample thickness, with a 40x and 60x oil objective with 200 ms exposure time, and 50 nW, 405 nm, 488 nm, 561 nm, and 640 nm laser exposure at 44% laser intensity. The imaging was performed from apical to basal surface so that peripodial cells were imaged first followed by the columnar cells of the wing disc. Optical slices were taken at distances equalling half the compartment length. Tiling was performed on images to get the entire sample in the field of view, and QuickStich^46^ was utilized to stitch individual tiles.

#### Image Analysis

For the Oregon-R staging data, CSBDeep^47^ was used on the Actin fluorescent channel for denoising (Fig. 1-6). For visualization purposes background subtraction of the pMyoII channel data was done using a rolling ball algorithm in ImageJ^48^. Intensity values for quantifications in this paper were measured using the raw data without any image corrections. Brightness and contrast adjustments were done to minimize saturation during the visualization of Rho1 data for InsR and tkv mutants (Fig. 7-8). Analysis of curvature at the basal surface of a *Drosophila* wing imaginal disc cross-section was carried out using an open-source package Kappa^49^ within ImageJ. StarDist^50^, an open-source ImageJ plugin was used to segment the nuclei. Details about quantifications of nuclear positioning and cell height can be found in the supplementary information of the text.

### Description of the computational model

#### General description of the model

The general Subcellular Element (SCE) modeling approach^33,51,52^ was adopted as the basis for the model presented here. This computational framework is used to simulate a two-dimensional (2D) cross-sectional profile of the *Drosophila* wing imaginal disc along the anterior-posterior (AP) axis. Using a 2D approximation allows for modeling large numbers of cells with high resolution and with special attention to mechanical cell properties and small changes in tissue structure and shape. Given that the cross-section is composed of boundary cells, squamous cells and columnar cells, the model includes a description for each cell type (Supplementary Fig. 10). Important distinctions from our previous model^33^ are the new model components presented in this paper which provide a detailed representation of cell division and actomyosin contractility in different parts of a cell. The model includes membrane nodes, nucleus nodes, and ECM nodes. Energy potentials are used to describe interactions between different nodes as in Nematbakhsh et al. 2020^33^. The total potential energy can be calculated across the whole model system and the position of each node of interest is determined via the following Langevin equations of motion:

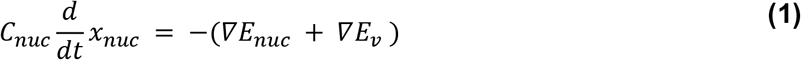

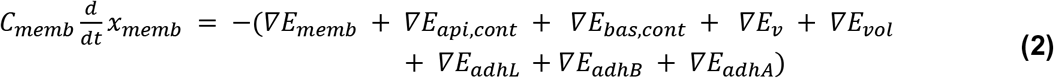

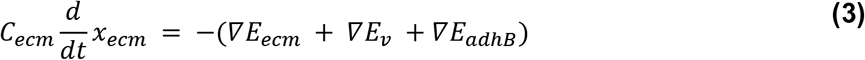

where *C*_*nuc*_, *C*_*memb*_ and *C*_*ecm*_ are damping coefficients. A linear spring potential, 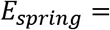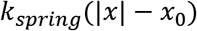 where *x*, *x*_0_ are the lengths between connected nodes and the resting length respectively, is used to describe the elasticity of the cell membrane (*E*_*mem*_) and the ECM (*E*_*ecm*_) (Supplementary Fig. 10A-D). Adhesion between membrane nodes residing in different cells (*E*_*adhL*_ and *E*_*adhA*_) and the membrane-to-ECM adhesion (*E*_*adhB*_) are also modeled via the linear spring potential (Supplementary Fig. 10D-F). Volume-exclusion (*E*_*v*_) between different types of nodes is described by the Morse potential. Since we introduce apical actomyosin contractility in this paper, an additional energy potential term is introduced in equation (2) and denoted by *E*_*api,cont*_. In this way, we capture actomyosin contractility in the apical (*E*_*api,cont*_) and basal (*E*_*bas,cont*_) parts of a columnar cell (Supplementary Fig. 10D). For completeness, we reproduced the table of energy potentials (Table 1) from our previous publication^33^, with the addition of the description of the new potential *E*_*api,cont*_, in order to provide a detailed description of the potential functions in the right-hand side of equations (1)-(3).

**Table 1.** Energy potentials used in the SCE model

| Potential function | Type | Biological representation |
| --- | --- | --- |
| $E_{nuc}$ | Morse potential | Size of a cell's nucleus |
| $E_v$ | Morse potential | Volume exclusion |
| $E_{memb}$ | Spring | Elasticity of the cell membrane |
| $E_{api,cont}$ | Spring | Apical actomyosin contractility inside the cells and above the nucleus |
| $E_{bas,cont}$ | Spring | Basal actomyosin contractility inside the cells and below the nucleus |
| $E_{vol}$ | Lagrange Multiplier | Volume conservation of the cytoplasm for each cell |
| $E_{adhL}$ | Spring | Cell-cell adhesion mediated by E-cadherin |
| $E_{adhB}$ | Spring | Cell-ECM adhesion mediated by Integrin |
| $E_{adhA}$ | Spring | Adhesion between columnar and squamous cells |
| $E_{ecm}$ | Spring | Extracellular matrix stiffness |

The use of the Langevin equation requires the assumption that cellular motion is under an overdamped regime which is valid in most of the micro-scale biological systems^33,51,53,54^. A detailed presentation of the types of potential energies and model construction can be found in Nematbakhsh and Levis, et al.^33^

#### Anisotropic apical and basal actomyosin contractility

A new feature of the current model provides a representation of anisotropic actomyosin contractility in the apical and basal parts of a cell. Namely, interactions between actin and myosin lead to the formation of a contractile force that creates constriction in the apical and basal portions of the columnar cells. In our computational model, this constriction effect is represented by the contractile springs linking the lateral sides of a columnar cell. Note that the interaction between actin and myosin is mostly membrane-bound and therefore the contractile force is exerted on the membrane surface altering the circumference of the cross-sections of each columnar cell.

While such contractile force may manifest in multiple directions, we only considered the apical and basal contractile forces that are perpendicular to the apical-basal direction, which constrict the lateral sides of the cells, as the dominant factor giving rise to the elongated rectangular geometry of the columnar cells. Therefore, in a 2D setting, this circumferential contractile force is projected on the 2D plane where our cross-sectional model tissue resides, in the form of linear springs. Thus, there is no direct interaction between the nuclei and the contractile springs, although some model simulations may falsely suggest otherwise. The contractile force exerted by the actomyosin network on the cell membrane is therefore described by the linear spring potential, 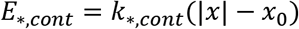 where *x* is the distance between a pair of membrane nodes connected by the contractile spring. A detailed description of the application for the anisotropic actomyosin patterning can be found in the following section as well as the last section.

#### Spatial representation of the model of the imaginal wing disc

The cross-sectional view of a wing disc can be roughly separated into left, middle, and right regions. The middle section is often found to be relatively flat in comparison to the left and right sections while significant bending is observed at the junctions between the three sections (Fig. 1). Based on this observation, we first simplified our assumption of the anisotropic basal and apical contractility patterns as a step function. For details of the computational model related to membrane stiffness and membrane-to-membrane adhesion, please see^33^. To systematically study the effect of perturbing both apical and basal actomyosin contractility, we divided the columnar cells of the model tissue into three uniformly spaced sections, for example, the apical contractility profile is set to be (*a*, 1.0, *a*) and the basal contractility profile is set as (1.0, *a*, 1.0) where 0 ≤ *a* ≤ 1.0 (Fig. 4). These manually chosen anisotropic patterns of actomyosin contractility are utilized to perform model validation. Such validation is necessary to confirm that the resulting physical behavior of the model is plausible and helps narrow down the range of possible perturbations to evaluate. These simulation results are in a good agreement with experimentally observed tissue shapes (Fig. 4E).

#### Cell growth and mitotic rounding

Several key features were developed to study the effect of cell growth and division on the shape generation of the wing disc. Such features include cell growth via the increase of cell volume, mitotic rounding events, and cell division. The volume conservation (or constraint) in the existing model utilizes a Lagrange multiplier energy function, 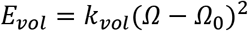 where 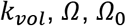 are the strength of the enforcement of volume constraint, current cell volume, and target (equilibrium) cell volume, respectively. To represent the increase in cell volume during cell growth, the equilibrium cell volume is increased linearly by a fixed value per simulation time step, namely 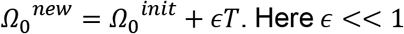 Here *ϵ* << 1 is a small positive real number representing the increment in volume per simulation time step, *T* is the current total simulation time, and 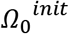 is the equilibrium cell volume at the starting time point of cell growth. Furthermore, the maximal value 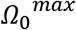 for 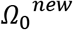 is set based on the expected cell volume before cell division occurs. Note that 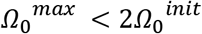 since the cross-section of the cell does not double even if the 3D volume is doubled.

### Modeling the basal constriction effect and the dynamics of actomyosin patterning during mitotic rounding

Mitotic rounding in the *Drosophila* wing disc is special in that the columnar cells experience an increased basal constriction that eventually pushes the majority of the cell interior content apically. This leads to an enlarged sphere at the apical side whose diameter is roughly a four-fold increase compared to the non-growing cell^51^. During this process, it was experimentally observed that the enrichment of actin and myosin concentration occurred near the basal section of the cell. This increased constriction effect is modeled as a time-dependent gradual increase in the number of actomyosin contractile springs in the existing model (Fig. 4) using the equation 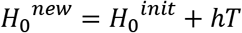. Here 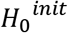 is the value at the basal point calculated as the average of the membrane node coordinates in the basal portion of the cell, *h* << 1 is a positive real number representing the increment per simulation time step in actomyosin network buildup, and *T* is the current total simulation time. *h* is approximated by using the experimentally observed mitotic rounding time duration and the portion of a mitotic rounding cell with high intensity of actomyosin presence at the onset of cell division (Fig. 1C). In this way, a contractile spring is active when the distances from the membrane nodes connected by such contractile spring to the basal point (*H*_*cont*_) satisfy 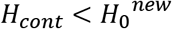. Therefore, as 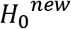 increases, the number of contractile springs also increases. The strength of the contractile springs is also increased to further increase the constriction effect. This combined effect would correspond to the increased actin and myosin concentration observed during the mitotic rounding process. Finally, at the end of the mitotic rounding process, we restore the value of the contractile spring constant to the initially prescribed value and assume that the actin concentration undergoes a restoration from a depolymerized state back to a pre-mitotic state.

#### Modeling cell division process

Cell division is modeled by constructing a division plane based on the centerline of each cell. Since the cell geometry can become rather distorted when simulating the mitotic rounding process, the division plane has to be carefully crafted. This was achieved by calculating the midpoints of the cell. These midpoints are based on the corresponding pair of nodes positioned at the two lateral sides of a given model cell. Given such a centerline, new cell membranes in the model can be constructed by shifting the centerline slightly toward the two lateral sides leading to two separated cells. This algorithm also corresponds to the scenario in that the planar orientation of the mitotic spindle is established as observed in the wild-type tissue, leading to the division along the lateral direction.

Each cell that can undergo cell growth and division was assigned an initial growth progress (*GP*) value *GP*_0_ < 1. Such value was increased by a fixed value *δ* << 1 per simulation time step. A cell in the model is considered to enter the mitotic rounding process when (*GP*_0_ + *δT*) > *GP*_*mit*_, where *T* is the current total simulation time, and *GP*_*mit*_ satisfying *GP*_0_ < *GP*_*mit*_ < 1 is a threshold value. When (*GP*_0_ + *δT*) ≥ 1, the cell is considered to reach the end of the M phase of the cell cycle leading to subsequence cell division. To ensure that premature cell division with respect to cell volume increase does not occur, the small increments used in the cell volume and actomyosin contractile spring are calibrated so that the cell growth and division process can complete in a timely manner. After a division event completes, the resulting cells are assigned a new *GP* value to indicate how fast such cells will undergo the mitotic rounding and cell division process again.

To ensure numerical stability in our simulations, the advancement in the cell cycle of a given cell is suspended if its immediate neighboring cell is already undergoing mitotic rounding and cell division. In terms of biological relevance, this is equivalent to the assumption that cells under high external stress are prevented from entering the mitotic phase. Lastly, in the simulation results presented in this paper, each cell can perform a maximal number of 10 cell divisions.

#### Modeling the introduction of a new cell in the cross-section

At the end of cell division, a random number is generated to determine whether a new cell is introduced into the same plane (or cross-section). Since cell division can produce a new cell that would lie within the same cross-section or outside of the cross-section, it is necessary to consider both in-plane and out-of-plane cell division. The probability of this event can be experimentally approximated by calculating the frequency of new cells occurring in the same cross-section post-division. If a new cell is not introduced in the same cross-section post cell division, the equilibrium volume is restored back to the original value 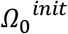 by a method discussed in an earlier section, except we use expression 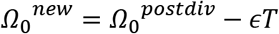 where 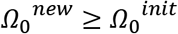.

Due to the unique mitotic rounding process, special care was taken to ensure numerical stability (Fig. 3). Since columnar cells in the model are tightly packed and experience an increase in cell volume during mitotic rounding, volume exclusion is manipulated to prevent membranes from different cells from overlapping. This is done via a temporary but significant increase in the magnitude of the volume exclusion energy potential coefficient which becomes active as a cell enters the mitotic process and briefly post cell division.

As described earlier, the number of contractile springs corresponds to the presence of actin filaments found throughout the wing disc. In this model, we use the actin intensity (or concentration) observed in experiments to determine the number of contractile springs per cell in the model. An additional simplification was applied by calculating the average value of the recorded intensity according to its corresponding spatial position on the wing disc (Fig. 2). However, this simplification can be discarded if the actomyosin intensity is governed by a time-dependent chemical signaling mechanism, and individual intensity can be prescribed to each individual cell.

#### A subcellular element model of nuclear and cell shape dynamics during organ growth

It still remains unclear how morphogenesis arises from the interplay between mechanical contraction and signaling networks. To fill this gap of knowledge, we specifically explore the relative contributions of proliferation and actomyosin contractility in controlling curvature, height, and nuclear positioning. During actomyosin contraction, pMyoII is responsible for bringing actin filaments closer to each other generating the contractile force. Such a force also depends on the availability of actin filaments^55^. The model incorporates actin density by utilizing a certain number of (linear) contractile springs in different parts of each cell. The impact of myosin on actin is represented by the spring constant of a contractile spring. In this paper, we consider both apical and basal contractility.

#### Determination of the number of springs in non-mitotic cells

The number of contractile springs within a non-mitotic cell depends on the height of the cell (*H*_*cell*_) and the local actin intensities. To determine the number of springs, we first need to determine the apical and basal portions of a cell that can be occupied by springs. Measuring from the apical side of a cell, the apical contractile spring height is denoted by 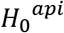, which defines the upper region of the cell where apical springs can exist (Fig. 2B, right panel, apical light blue region). Similarly, measuring from the basal point of a cell, we have the basal contractile spring height 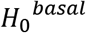(Fig. 2B, right panel, basal dark blue region). The exact values of 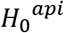 and 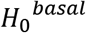 at the current iteration are determined by the local actin intensities. More specifically, we assume that 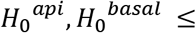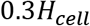 and define 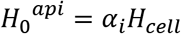 and 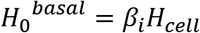 The weight constant *α*_*i*_ is defined by *α*_*i*_ = 0.3(*A*_*i*_/*A*_*max*_) where *A*_*i*_ and *A*_*max*_ represent the local apical actin intensities and the maximum apical actin intensity, respectively. The calculation for *β*_*i*_ is similar to *α*_*i*_, except we utilize the local and maximum basal actin intensities. Finally, a contractile spring becomes active if the distance between the apical point of the cell and the nodes connected by the contractile spring is less than 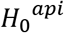 or greater than 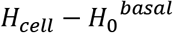. If this distance is less than 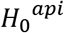,an apical contractile spring will manifest in the upper portion of the cell while if the distance is greater than 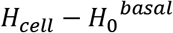, then a basal contractile spring becomes active.

## Supporting information

Supplementary Information: Balancing competing effects of tissue growth and cytoskeletal regulation during Drosophila wing disc development

## Code availability

An open-source C++ and CUDA implementation of the computational model is available on GitHub (https://github.com/librastar1985/Episcale_CrossSectionalView) or in Zenodo (https://doi.org/10.5281/zenodo.7087728)^56^.

## Acknowledgments

This work was supported by the National Science Foundation grant NSF2029814 to all co-authors and by a pilot grant from the Northwestern University NSF-Simons Center for Quantitative Biology to JZ. The authors would also like to thank members of the Zartman lab for helpful discussions. Model simulations were run on UC Riverside HPCC.

## Contributions

N.K. prepared the original draft. N.K. and M.S.M. performed experiments, and N.K. analyzed the data. K.T. developed and calibrated the computational model, performed simulations, and wrote the manuscript. M.S.M. and W.C. contributed to the data and simulation analysis, and edited the manuscript. J.R.A. performed simulations, contributed to the data and simulation analysis, and edited the manuscript. J.Z. and M.A. conceived, designed, analyzed and interpreted results, supervised this study, and wrote the manuscript. All the authors read and agreed to the manuscript.

