## Supplementary Information: Balancing competing effects of tissue growth and cytoskeletal regulation during Drosophila wing disc development for "Balancing competing effects of tissue growth and cytoskeletal regulation during *Drosophila* wing disc development"

### Contents

S1. Supplemental details of methods and image analysis
S2. Results and supplementary figures
  S2.1. Proliferation decreases with pouch size.
  S2.2. Collagen IV as a key structural component of the ECM. Integrins are dominantly localized at the anterior and posterior end of the basal surface as compared to the center of the pouch.
  S2.3. Integrins and Collagen IV co-localize with Actin within the cortical region of a dividing cell.
  S2.4. ECM stiffness plays a minor role in regulating the anisotropic basal curvature and cell height compared to actomyosin contractility.
  S2.5. The local height of tissue is regulated by differences in apical-basal contractility and specification of cell volume.
  S2.6. Activation of Dpp signaling initiates elongation of pouch cells.
  S2.7. Inhibiting Dpp signaling activity downregulates Rho1 expression and reduces inwards bending at the lateral domains of the pouch.
  S2.8. Effect of medial tissue flattening on tissue pressure and membrane tension.
S3. Computational modeling overview
S4. Image analysis and data quantification pipelines
S5. Supplementary videos
S6. Tables
S7. SI References

### S1. Supplemental details of methods and image analysis

**Additional method details for Fig. 1.** *Drosophila* wing imaginal discs belonging to different stages of larval development were dissected and fixed with Phalloidin and DAPI to mark Actin and nuclei, respectively (Fig. 1A). Further, an antibody staining against phosphorylated non-muscle Myosin II (pMyoII) was carried out to measure the spatiotemporal dynamics of myosin within the developing epithelia. Cross-sections along the DV axis were analyzed to study the role of Dpp signaling during development (Fig. 1E). Kappa, an open source FIJI plugin, was used to quantify the local curvature of basal epithelia for the DV optical section[1]. The calculated local curvature has been color-coded and overlaid on top of the basal surface of the pouch as shown in Fig. 1B. A custom MATLAB-based pipeline (Supplementary S4, Supplementary Fig. 11) was then used to calculate the thickness of the tissue and the positioning of nuclei with respect to the basal surface along the DV axis (Fig. 1C, Fig. 1D).

**Additional method details for Fig. 4.** For model calibration, wing imaginal discs belonging to 96 h AEL were dissected and fixed with Phalloidin and DAPI to mark the cell boundaries and nuclei respectively (Fig. 4A). Kappa, an open-source package in ImageJ, was used to quantify the curvature of the basal epithelia for DV optical sections[1] (Fig. 4A'). Points on the basal surface were selected and used to fit a spline in order to define the basal surface in 2D. Absolute values of curvatures at uniformly sampled points on the fit basal surface were plotted against the normalized distances from the AP axis for a total of six samples (Fig. 4B). Distances of points from the AP axis measured along the basal surface were normalized against the total basal curve length. In addition to curvature, Kappa was also used to quantify the fluorescence intensity of pMyoII across the apical and basal surfaces along the DV axis. The location of these points in the apical and basal surface along the DV axis was normalized against the total length of the apical and basal surface respectively. The fluorescence intensity was normalized further against the maximum reported intensity value across both surfaces and plotted as a scatter plot against the normalized location of points on the corresponding surfaces on the DV axis. Further, the apical and basal surfaces were subdivided into three equal domains based on the respective curve length and the averaged fluorescence intensity in the respective domain has been plotted as a solid line (Fig. 4D).

**Additional method details for Fig. 5.** In this section, we quantified the correlations between concentration levels of pMyoII on the apical and basal surfaces of cells and morphological metrics of wing imaginal disc such as local tissue thickness and basal epithelial curvature. The DV section of the wing imaginal disc was split into three equal domains using the total length of the basal surface for sample discs belonging to different stages of development. Next, we analyzed Kappa[1]-estimated local curvature values at several points sampled on the basal surface of the medial pouch domain. The patterning of local curvature across the medial basal domain has been represented as a kernel density estimate where the y-axis represents the probability distribution function for the local curvature values represented along the x-axis[2]. The kernel density estimates of the medial-basal local curvature for discs belonging to different ages are stacked vertically to create a ridgeline plot (Fig. 5C). The kernel density estimates are further color-coded based on the median of the local curvature values highlighting a decrease in medial basal curvature with the age of disc. A custom MATLAB-based pipeline was then used to discretize the DV section of the pouch into a total of 90 elements each theoretically mimicking a pouch columnar subtype cell as shown in Supplementary Fig. 12 (Supplementary S4). For each discretized cell, local tissue thickness along with median expression of pMyoII across the apical and basal surface was calculated. This information was used to calculate the ratio of apical to basal relative concentration levels of pMyoII ($pMyoII_{apical}$ / $pMyoII_{basal}$). Next, kernel density estimates[2] of $pMyoII_{apical}$ / $pMyoII_{basal}$ in cells within the medial and lateral domains were approximated using MATLAB for samples belonging to different stages of development. Individual estimates were stacked vertically to create a ridgeline plot (Fig. 5D). Through the plots, it can be clearly seen that there is an increase in apical to basal ratio of pMyoII with the age of the disc at the medial pouch domain.

We also used the image analysis pipeline on discs belonging to different stages of development for quantifying correlations between the ratio of $pMyoII_{apical}$ / $pMyoII_{basal}$ and local tissue height. A 2D scatter plot was first plotted with the x-axis and y-axis representing $pMyoII_{apical}$ / $pMyoII_{basal}$ and local tissue height, respectively (Fig. 5G). Data extracted from each sample was plotted using unique colors. First, linear regression models were fit individually to points extracted from each sample using the *fitlm* function of MATLAB. The predictions made by the model are represented by a solid color-coded line. Next, a single linear model was fit to the aggregated data from different samples represented by a solid black colored line. Averaged $R^2$ values for samplewise model fits along with the $R^2$ value of the global model fit have been reported. A p-value for an F-test was calculated to evaluate the statistical significance of the global fit. Plotting and statistical tests were carried out in MATLAB. We also report the presence of a gradient in tissue height across the DV axis. The tissue thickness is maximum at the pouch center and decreases gradually as one moves towards the lateral sides (Fig. 5H). For samples at a particular developmental stage, the variation in tissue thickness profile is first plotted as a transparent solid colored line. Next, we used the *fit* function in MATLAB to fit a gaussian equation to model the variation in cell height. Model predictions have been plotted as a solid opaque colored line. Finally, a single gaussian equation was fit to the combined data from all samples. The width at half maximum of the gaussian fits (WHM) was also estimated for the local and global gaussian models in order to estimate the decay lengths of the tissue thickness profile.

**Additional method details for Fig. 6.** DV optical sections for wing imaginal discs belonging to 72 h and 96 h AEL larval stages were analyzed. The proximity of nuclei with respect to the basal surface has been defined as the ratio of the distance of nuclei from the basal surface ($d_B$) over the sum of the distances of nuclei from apical and basal surfaces ($d_A$, $d_B$). Details about the quantification pipeline are mentioned in supplementary S4 (Supplementary Fig. 11). As described in previous sections, the cross-section was subdivided into three regions i.e. one medial and two lateral regions. Nuclei in each subregion were identified and their proximity to the basal surface was calculated. A total of 5 samples from 72 h AEL and 10 samples from 96 AEL were analyzed. The proximity of nuclei from the basal surface for nuclei belonging to the medial and lateral subdomains for all the samples from the two developmental stages have been plotted as a combination of beeswarm and violin plots using the ai_goodplot[3] function in MATLAB (Fig. 6D). A two-sample t-test was carried out to measure the statistical significance and the corresponding p-values have been reported.

**Additional method details for Fig. 7.** GAL4/UAS[4] system was used to express dominant recessive and constitutively active form of insulin receptors (InsR) in the posterior compartment of wing imaginal disc using an engrailed-Gal4 driver (Fig. 7D, E). The engrailed-Gal4 also expresses a UAS tagged with a GFP marker to fluorescently label the posterior half, i.e., the region of perturbation. Wing imaginal discs belonging to 100-120 h AEL (wandering larvae, early 3$^{rd}$ instar) were dissected and fixed. Samples were treated with DAPI to label nuclei. Additional Rho1 antibody staining was carried out to label cell and tissue boundaries for tracking morphological changes within the wing imaginal disc. An Anti-Phospho-Histone H3 (PH3) antibody staining was carried out to mark the mitotic cells within the pouch.

**Additional method details for Fig. 8.** GAL4/UAS[4] system was used to express the constitutively active form of Dpp receptors, Thickveins (Tkv), in the posterior compartment of wing imaginal disc using an engrailed-Gal4 driver (Fig. 8A-C). Samples dissected from 100-120 h AEL were dissected and fixed. DAPI was used to label the nuclei. A PMAD antibody staining was performed to validate the UAS-Tkv$^{CA}$ line (Fig. 8A'-C'). A Rho1 antibody staining was also carried out to measure changes in tissue geometry. Rho1 is also upstream of pMyoII and hence can be used as a proxy to measure contractility (Fig. 8A''-C'').

### S2. Results

**S2.1 Proliferation decreases with pouch size.** In this section, we report a a decrease in cell proliferation with an increase in age of the disc . The obtained result is in accordance with previous literature studies[5]. Wing imaginal discs were dissected from larvae of different sizes belonging to different stages of development. The dissected discs were fixed and an anti-PH3 antibody staining was carried out to label the proliferating cells. The mitotic index was next defined as the ratio of the area of cells that are marked by PH3 over the area of the pouch (Supplementary Fig. 1A). A 2D scatter plot was plotted with the X-axis representing the pouch area and the Y-axis the mitotic index (Supplementary Fig. 1B). An exponential decay model was next fit to the data using MATLABs fit function.. The model details along with the goodness of fit indicated by a R2 value has been included as a plot inset. The tendency of cells to divide decreases as the size of the pouch increases.

**S2.2 Collagen IV as a key structural component of the ECM. Integrins are dominantly localized at the anterior and posterior end of the basal surface as compared to the center of the pouch.** Wing imaginal discs collected from early 3rd instar larval disc of Oregon-R were fixed and antibody staining was performed to visualize the localization of Collagen IV and Integrins. Collagen IV is one of the structural components of ECM while Integrins facilitate adhesion of basal cell surfaces with the ECM to facilitate extracellular signaling. Optical sections along the DV boundary were analyzed (Supplementary Fig. 2A). Both Collagen IV and Integrins are primarily localized at the basal surface with a diminished expression on the apical side of the columnar cell (Supplementary Fig. 2A-B). Further, the localization in the basal surface of the pouch cell is increased at the region with higher local curvature, i.e., either towards the anterior or posterior direction (Supplementary Fig. 2A'-B'). This provides evidence of the existence of anisotropy in Integrin-mediated ECM adhesion.

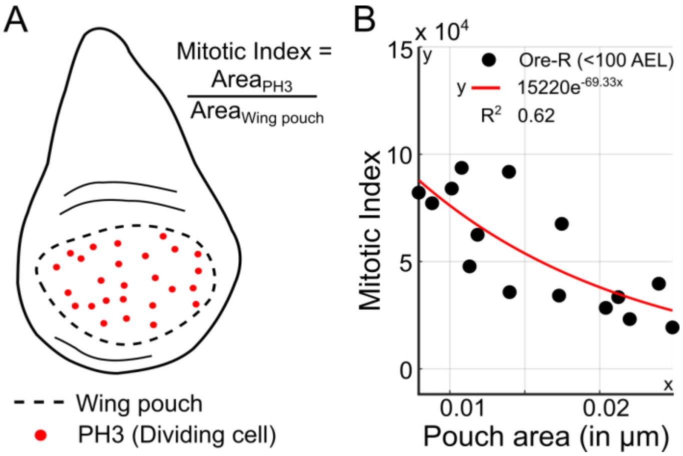


**Supplementary Figure 1. Cell proliferation decreases with the age of the pouch. (A)** Mitotic index is defined as the ratio of the area acquired by cells expressing Phospho Histone-H3 (PH3) over the area of the wing disc pouch. **(B)** Plot showing the variation of the mitotic index with respect to the pouch size. The red line indicates an exponential fit obtained through regression in MATLAB...

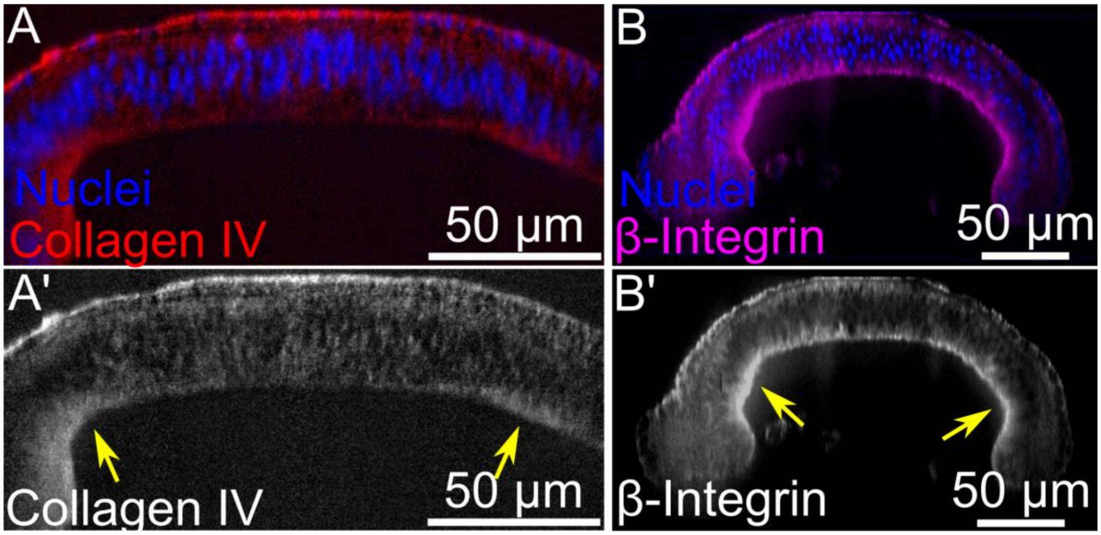


**Supplementary Figure 2. Collagen IV and Integrins are basally localized predominantly at regions with high local curvature. (A-A')** DV cross-section of a wing imaginal disc dissected at ~100h AEL. Fluorescent labels indicate nuclei and localization of Collagen IV, a key structural component of ECM. **(B-B')** DV cross-section of a wing imaginal disc dissected at 96 h AEL. Fluorescent labels indicate nuclei and localization of $\beta$-Integrin.

**S2.3 Integrin and Collagen IV co-localize with Actin within the cortical region of a dividing cell.** Wing imaginal discs belonging to the early 3rd instar larval stage were dissected and fixed. DAPI and Phalloidin were used to label the nuclei and Actin, respectively. An antibody staining against anti-Collagen IV and anti-Integrin was carried out to visualize spatial patterning of Collagen IV and Integrins within the fixed tissue. Based on the data from Actin and nuclei fluorescence channels, a mitotic nucleus was selected within the pouch (Supplementary Fig. 3A). An optical slice along the long axis of the dividing cell was taken to have a lateral perspective of the dividing cell (Supplementary Fig. 3B). A major highlight of a dividing cell is the presence of higher levels of Actin around the cortical region of a dividing cell[5]. It can be seen clearly from the data that along with Actin, Integrin and Collagen IV also colocalize at the cortical region of the dividing cell.

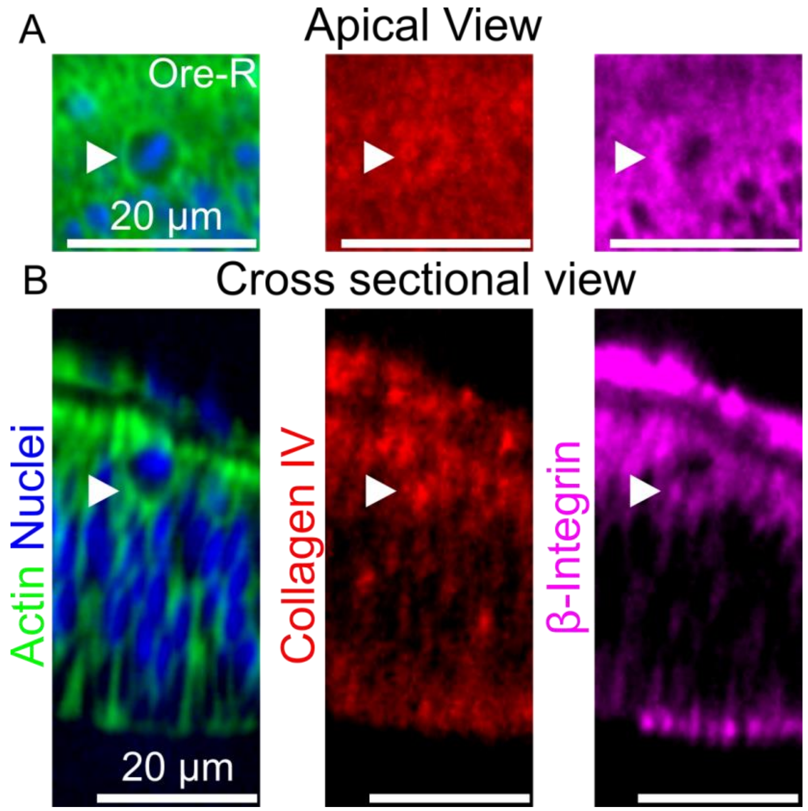


**Supplementary Figure 3. Localization of both Collagen IV and Integrin around a dividing cell. (A)** Apical view of a dividing cell. **(B)** Cross-sectional view of a dividing cell.

**S2.4 ECM stiffness plays a minor role in regulating the anisotropic basal curvature and cell height in comparison to actomyosin contractility.** Simulations were carried out to evaluate the role of ECM stiffness on the tissue shape and columnar cell heights. We set the anisotropic actomyosin profile as presented in Fig. 4D. When comparing the simulation with unchanged basal ECM stiffness ($k_{ecmc}$) (Supplementary Fig. 4A) to the case where the basal ECM stiffness is reduced by 93.75%, (Supplementary Fig. 4A') the resulting tissue shapes and cell heights remain marginally different (Supplementary Fig. B,C). Consistent with experimental observations, the anisotropicity of the tissue shape is dominantly determined by actomyosin contractility.

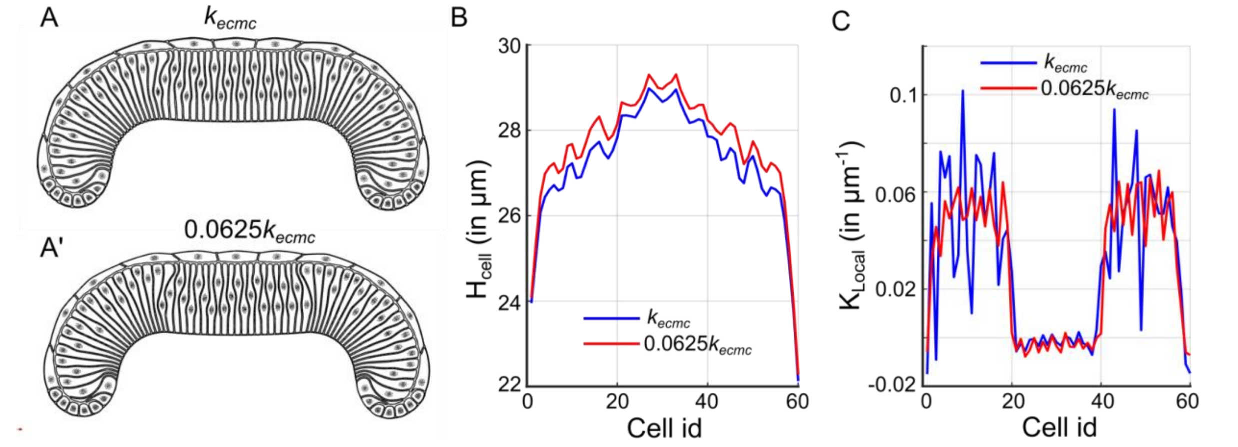


**Supplementary Figure 4. ECM stiffness plays a minor role in regulating the anisotropic basal curvature and cell height in comparison to actomyosin contractility. (A)** Unchanged $k_{ecmc}$.**(A')** $k_{ecmc}$ reduced by 93.75% **(B)** Columnar cell heights with reduced and unchanged $k_{ecmc}$.**(C)** Local curvature of basal epithelia with reduced and unchanged $k_{ecmc}$.

**S2.5 Tissue local height is regulated by the difference in apical-basal contractility and specification of cell volume.** To test if cell height is driven by the apical-basal stiffness of cells, we ran simulations with a differential patterning of contractility parameters across the apical and basal surfaces. The pouch was subdivided into 5 nested domains namely: left (L), left medial (LM), medial (M), right medial (RM), and right (R). Perturbation of $k_{api,cont}$ and $k_{bas,cont}$ was done only in the medial domain of the pouch. The parameter values for the three different cases are listed in a table in Supplementary Fig. 5B. Initially, we assumed the contractility to be localized only on the basal surface. We first increased both $k_{api,cont}$ and $k_{bas,cont}$ without changing the ratio of $k_{api,cont}$ / $k_{bas,cont}$. Analysis of the height profiles showed that an increase in levels of contractilities in the apical and basal surface without changing the ratio of the two increased cell height (Supplementary Fig. 5A', 5C). Next, we increased both $k_{api,cont}$ and $k_{bas,cont}$ in such a way that the increment also increased the ratio between the two. A slight change in height was observed as compared to the previous case (Supplementary Fig. 5A'', 5C).

In the next study, the volume of cells in the medial domain of the pouch was varied keeping the contractility parameters fixed (Supplementary Fig. 6). The volumes of cells in the pouch central region were increased by a factor of 10% and 20%. Increasing the volumes led to an increase in columnar cell height as well (Supplementary Fig. 6B). The parameter values used in the simulation are listed in Supplementary Fig. 6A. In summary, our results show that both cell volumes and contractility parameters (apical and basal) regulate columnar cell height within the tissue.

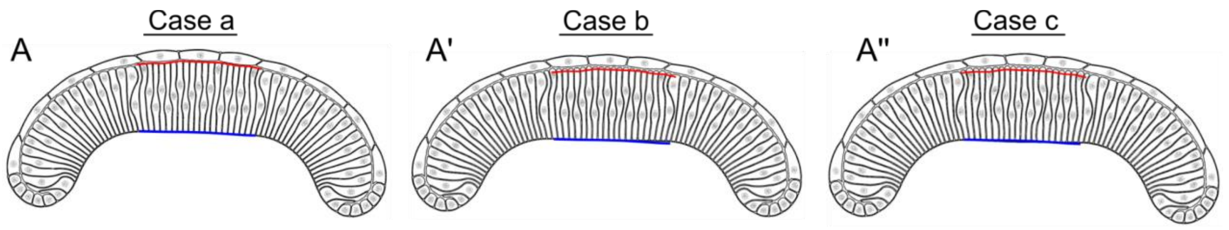


B

Contractility multiplier profile

| | | L | LM | M | RM | R |
|---|---|---|---|---|---|---|
| a | apical | 0.06 | 1.25 | **1.25** | 1.25 | 0.06 |
| | basal | 1.5 | 1.5 | **1.88** | 1.5 | 1.5 |
| b | apical | 0.06 | 1.25 | **1.56** | 1.25 | 0.06 |
| | basal | 1.5 | 1.5 | **2.4** | 1.5 | 1.5 |
| c | apical | 0.06 | 1.25 | **2.19** | 1.25 | 0.06 |
| | basal | 1.5 | 1.5 | **2.62** | 1.5 | 1.5 |

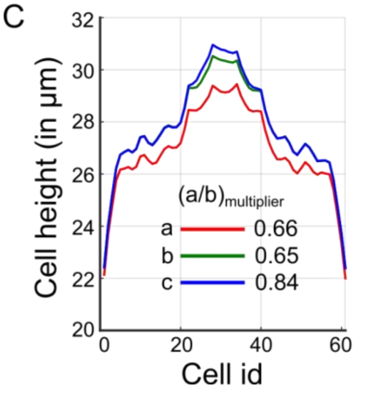


**Supplementary Figure 5. Effect of apical-basal contractility profiles on cell height. (A-A")** Stable shapes obtained as a result of varying actomyosin contractility in the pouch medial domain. **(B)** A table of parameters for the simulations run in this case study. **(C)** Height profiles in the tissue are plotted against the cell id (from left to right of tissue) for the different cases run in this study.

A

Contractility multiplier profile

| | L | LM | M | RM | R |
|---|---|---|---|---|---|
| apical | 0.06 | 1.25 | 2.19 | 1.25 | 0.06 |
| basal | 1.5 | 1.5 | 2.62 | 1.5 | 1.5 |

Volume multiplier profile

| | L | LM | M | RM | R |
|---|---|---|---|---|---|
| Case a | 1.0 | 1.0 | **1.0** | 1.0 | 1.0 |
| Case b | 1.0 | 1.0 | **1.1** | 1.0 | 1.0 |
| Case c | 1.0 | 1.0 | **1.2** | 1.0 | 1.0 |

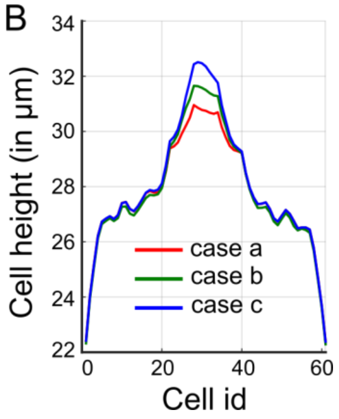


**Supplementary Figure 6. Effect of cell volume on cell height. (A)** A table of parameters for the simulations run in this case study. **(B)** Height profiles in the tissue are plotted against the cell id (from left to right of tissue) for the different cases run in this study

**S2.6 Activation of Dpp signaling initiates elongation of pouch cells.** GAL4/UAS system was used to express the constitutively active form of Tkv receptors in the posterior compartment of the wing disc using an engrailed-Gal4 driver (Supplementary Fig. 7A). An antibody staining against PMAD was carried out to validate the genetics of the cross-section. Higher fluorescence of PMAD in the posterior compartment suggests activation of the Dpp signaling pathway upon overexpression of $Tkv^{CA}$ (Supplementary Fig. 7A’). An anti-Rho1 antibody staining was also carried out to measure cell shape changes occurring as a result of the partial mutation in the posterior compartment (Supplementary Fig. 7A''). A 50 µm long cross-section perpendicular to the AP boundary indicated by the yellow dashed line in Supplementary Fig. 7A'' was taken across the dorsal compartment such that equal portions lie in the control (anterior) and $Tkv^{CA}$ expressing (posterior) domains of the pouch. Visualizing the reslice reveals thickening of the pouch near the AP boundary, localized primary at the central region of perturbation half (Supplementary Fig. 7B). Median tissue height in the control and perturbed domains was next calculated across the cross-sections from six similarly staged samples and plotted as a box plot in MATLAB. It can be seen that there was a ~6.3% increase in average tissue height. A two-sample t-test was next carried out to measure the statistical significance of this comparison[6]. A p-value of 0.07 has been reported.

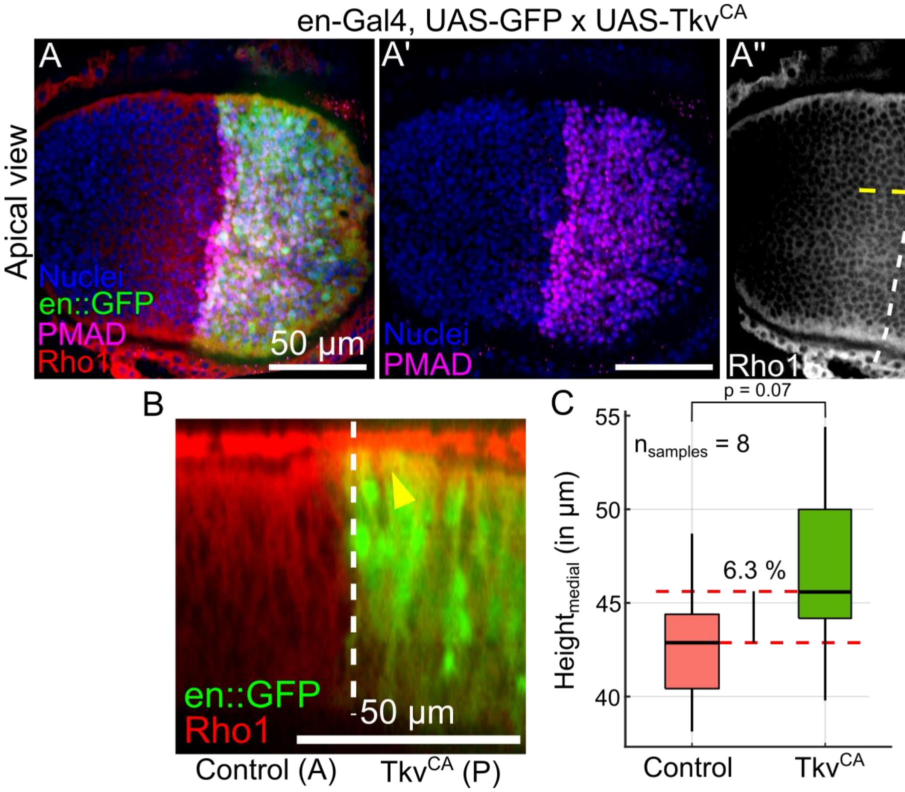


**Supplementary Figure 7. Expression of $Tkv^{CA}$ in the posterior compartment increases cell height. (A-A'')** Apical view of a wing disc expressing $Tkv^{CA}$ in the posterior compartment of the wing imaginal disc. Fluorescence labels have been indicated on the button left portion. **(B)** A cross-section along the yellow dashed line A'' is visualized. **(C)** Box plot showing the distribution of tissue heights across control and $TkV^{CA}$ expressing medial domains of the pouch.

**S2.7 Inhibiting Dpp signaling activity downregulates Rho1 expression and reduces inwards bending at the pouch lateral domains.** GAL4/UAS system was used to downregulate the activity of Dpp signaling in the posterior compartment of the wing disc using an engrailed-Gal4 driver and a commercially available UAS-$Tkv^{RNAi}$ line and DV cross sections of the discs were analyzed (Supplementary Fig. 8B). An anti-PMAD antibody staining was carried out for validation. It can be seen that expression of $Tkv^{RNAi}$ leads to a reduction in fluorescence intensity of PMAD signals in the posterior half as compared to the anterior half (Supplementary Fig. 8B'). Comparing this with the control, a wing imaginal disc for the parental engrailed-Gal4 driver, the peaks of fluorescence in both the anterior and posterior half are roughly comparable (Supplementary Fig. 8A'). Next, an anti-Rho1 antibody staining was carried out to measure changes in cytoskeletal regulation (Supplementary Fig. 8B''). Rho1 expression is known to promote the phosphorylation of myosin and hence is a regulator of tissue contractility[7]. We report a decrease in expression of Rho1 in the posterior $Tkv^{RNAi}$ expressing compartment as compared to its control anterior half (Supplementary Fig. 8B''). On the other hand, the expression of Rho1 for the control is symmetric for the anterior and posterior halves (Supplementary Fig. 8A''). Lastly, we also report a loss in inwards bendings at the lateral end of basal epithelia in the posterior half where the Tkv receptors were inhibited (Supplementary Fig. 8B).

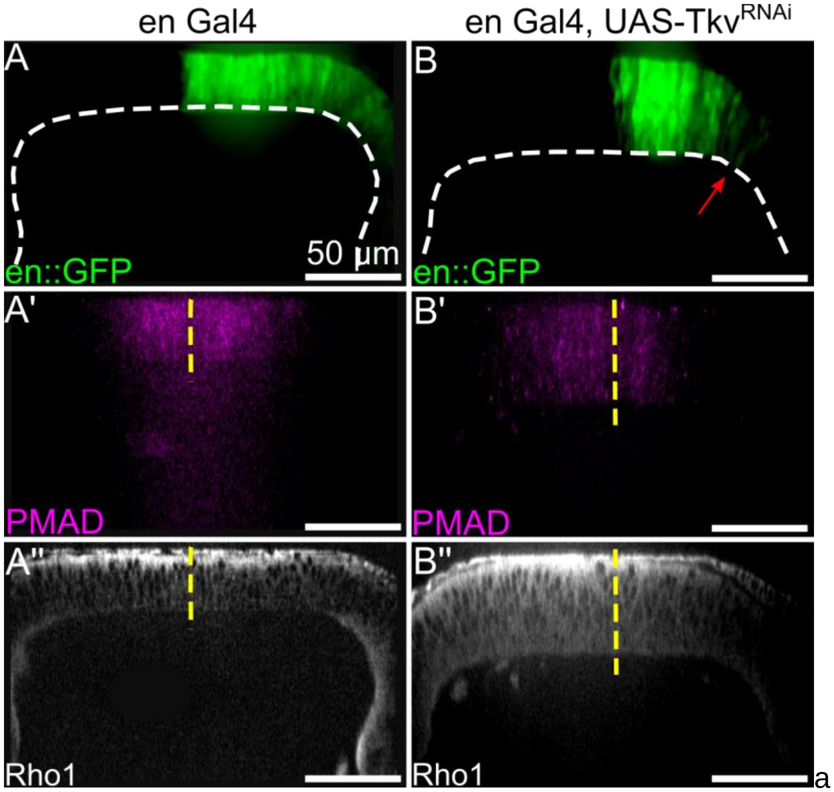


**Supplementary Figure 8. Inhibition of Tkv receptors results in a reduction of Rho1 followed by a loss in inwards bending of lateral pouch domains. (A-A'')** Cross section along the DV boundary of a control wing imaginal disc dissected from larvae of an engrailed-Gal4 driver. Fluorescence labels have been indicated in the lower left panel of each figure. **(B-B'')** Cross section along the DV boundary of a wing imaginal disc expressing $Tkv^{RNAi}$ in the posterior compartment. The posterior compartment also expresses GFP as indicated in A and B.

**S2.8 Effect of medial tissue flattening on tissue pressure and membrane tension.** "Stress" contains the normal force pushing against the cell membrane (Supplementary Fig. 9B), and it is calculated as the sum of the normal forces:

$$f_n = -2k_{area}(area\ cell_{current} - area\ cell_{equi})\sqrt{(posYR - posYL)^2 + (-posXR + posXL)^2},$$

where {posYR, posYL, posXR, posXL} are the x- and y-components of the vectors from the cell center to nodes on a membrane spring. It is then combined with the repulsion force of nuclei acting on the membrane nodes and divided by the cell perimeter length. Essentially, it represents the force acting on the membrane node from internal factors. "Membrane Tension" calculates the total force acting on a membrane node due to the stretching of linear springs connected to such a membrane node (Supplementary Fig. 9A). If the linear spring is below a minimum value, 1e-8, no tension force will be calculated. The value is calculated as: $k_{spring}(L_{current} - L_{equi})$

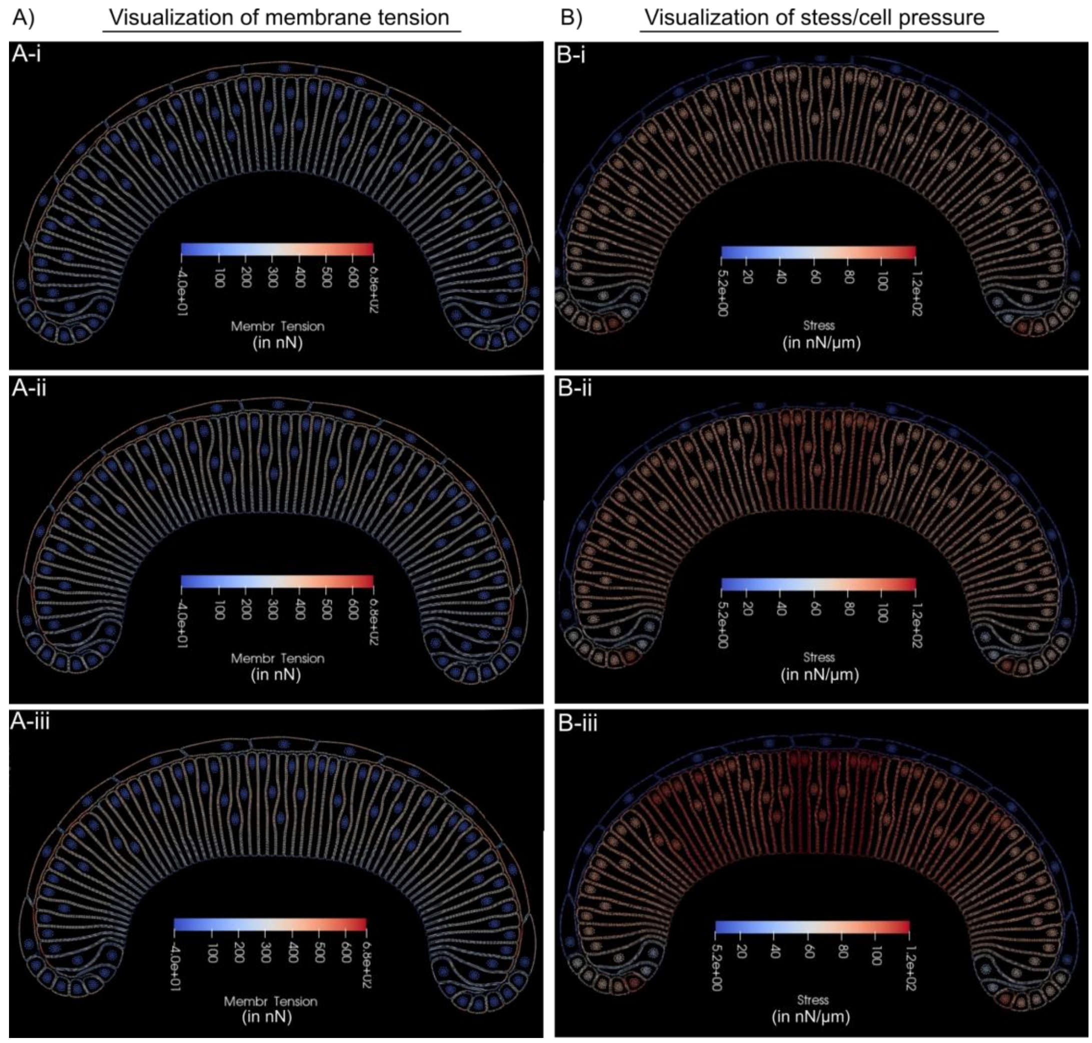


**Supplementary Figure 9. Color-coded visualization depicting: (A)** membrane tension and **(B)** stress/pressure

### S3. Computational modeling overview

We developed a multi-scale computational model that allows us to study how changes in local cellular mechanical properties govern the dynamical changes in epithelial tissue geometry. The subcellular element method provides unparalleled resolution in describing detailed intra- and inter-cellular interactions including cell-cell adhesion and membrane-nucleus interactions

(Supplementary Fig. 10). Previously, our subcellular element model was developed to uncover the novel role of ECM in maintaining the characteristic cross-sectional shape of the wing disc, which exhibits a “dome” -like structure[9]. Experimentally-obtained data such as the local actin and myosin intensities were used for model calibration.

Several previous computational models have been developed to study the impact of changing mechanical properties of epithelial tissue on its structure and shape[21–23]. A common setup in many previous models is to describe the epithelial layer as a tightly connected two- or three-dimensional structure composed of polyhedrons. As surveyed by Smallwood, earlier models can be traced back to the paper by Honda that describes the epithelial structure from a top-down view using Dirichlet domains[24,25]. Subsequent models, either two-dimensional or three-dimensional, were developed utilizing geometrical techniques represented by topology dynamics, center dynamics, boundary dynamics, or vertex-based dynamics where an individual cell is depicted as a polyhedron [24,26–33].

Physics-based models describing an epithelial cell layer where mechanical stress and elastic deformation are modeled via potential terms, stress-tensor, or continuum approach, are a key area of continued interest[26,34–37]. These computational models have been used to provide insight into important biological questions including the formation of epithelial folding and invagination in order to identify potential mechanisms driving such processes. For instance, in the work of Marin-Riera et al., an off-lattice center dynamics model was developed to illustrate the ability to capture the interaction between an epithelial layer with its surrounding extracellular matrix (ECM), and how differential adhesion, cell migration, and cell contraction lead to tissue deformation[36]. A key aspect of these models is allowing different degrees of contraction at the apical and basal surface, regardless of the model dimension, to achieve deformation. However, such models often make simplified assumptions including but not limited to reducing the contacting surfaces between two neighboring cells into a single surface, and therefore reducing the flexibility of the lateral cell surfaces[38–41]. In addition, it is also frequently assumed that the contraction is surface-bound, either apically or basally.

Recently, Tozluoglu et al. developed a 3-dimensional finite element model where each individual cell in the *Drosophila* wing pouch is represented as a triangular prism, and growth is modeled by the increase in the volume of prisms, to study the mechanism for fold formation[8]. Using this model capable of capturing the ability of cells to resist deformation including shear strain, they conclude that external resistance to growth from the ECM is essential for buckling the tissue and the increased apical stiffness can induce the correct number of folds. Ioannou et al. developed a three-dimensional hybrid vertex model that allows a more complex polyhedron representation of the epithelial cells improving the ability to more realistically capture the packing of cells[42]. The subcellular element method has also been utilized to study both the top-down and cross-sectional view of the epithelial layer in the *Drosophila* wing disc[43,44]. While more computationally intensive, the subcellular element approach enables a multiscale investigation including, but not limited to the interplay between cellular deformation due to contraction and the dynamics of subcellular components including the nucleus, which is often absent in more coarse-grained models.

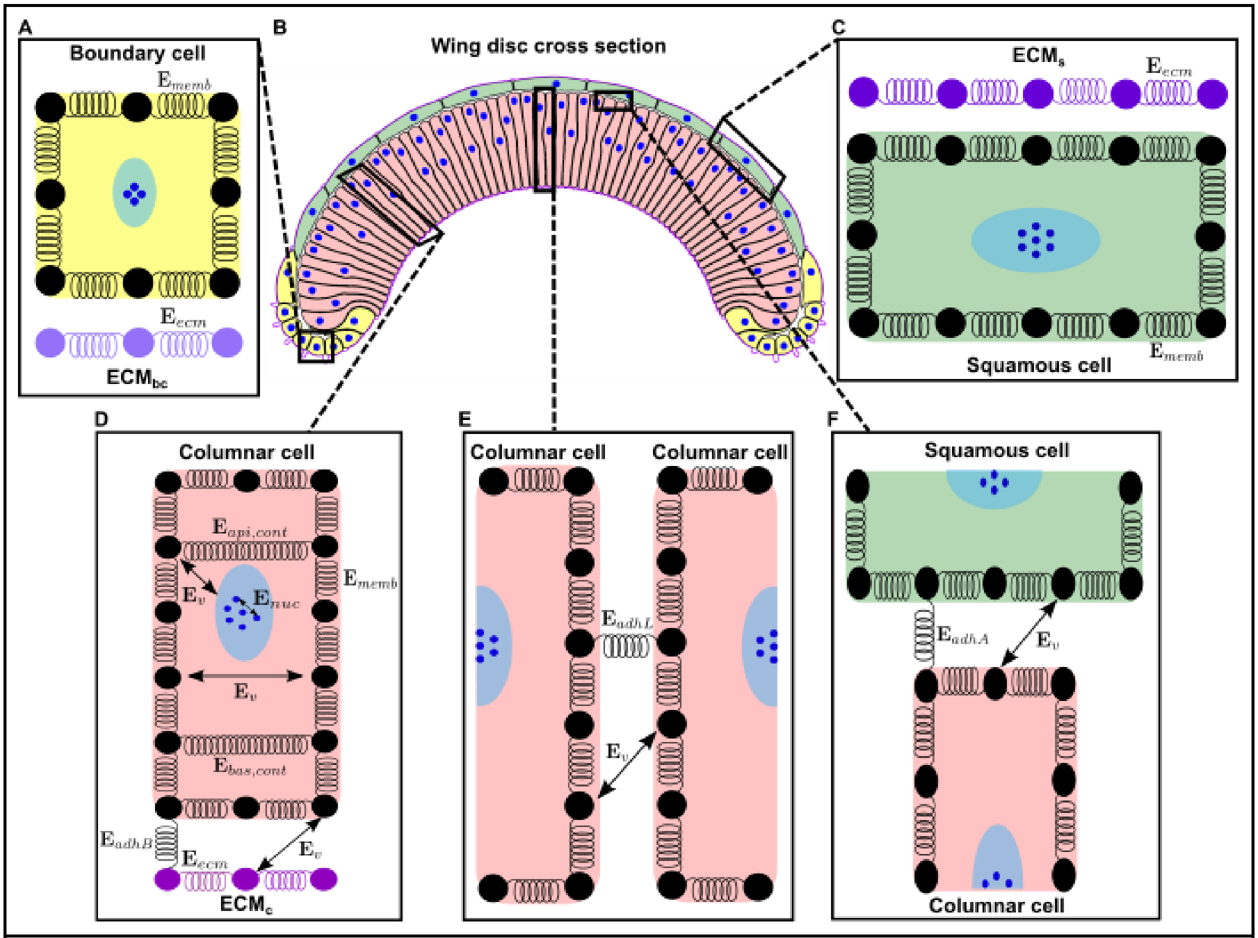


**Supplementary Figure 10. Two-dimensional (2D) multi-scale subcellular element (SCE) model of the *Drosophila* wing disc cross section along the anterior-posterior (AP) axis.** This model closely resembles the wing disc cross section by including a representation of the three different cell types (boundary, squamous and columnar cells) that make up the cross section and of the ECM which connects to the basal surface of the cells. Simulated cells, nuclei and ECM are represented by a set of nodes interacting via different potential energy functions. **(A)** Model representation of a boundary cell and its associated ECM denoted by $ECM_{bc}$. **(B)** Model simulation of the cross-sectional profile of the wing disc along the AP axis includes boundary cells (colored in yellow), columnar cells (colored in light red), squamous cells (colored in green) and the ECM (colored in purple). **(C)** Model representation of a squamous cell and its respective ECM denoted by $ECM_s$. **(D)** Model representation of a columnar cell with different potential energy functions that capture intracellular interactions, apical and basal actomyosin contractility and cell interactions with the ECM (denoted by $ECM_c$). **(E)** Adjacent columnar cell nodes interact via linear spring $E_{memb}$ and Morse potentials $E_v$ to maintain the cells closely connected without the membranes overlapping. **(F)** The apical membrane of columnar cells is connected to squamous cells.

There have also been models developed focusing on identifying the link between mechanical behaviors with the underlying chemical signaling pathways. The work by Hughes et al. has delved into how mechanical compaction of the extracellular matrix during mesenchymal condensation leads to tissue folding[45]. In the work of Zmurchok et al., a two-way feedback between signaling and mechanical tension which leads to a spectrum of cell contraction and relaxation was investigated to observe waves of contraction and relaxation sweeping through a two-dimensional model epithelium[46]. However, it still remains unclear how morphogenesis arises from the interplay between mechanical contraction and signaling networks. To fill this gap of knowledge, we specifically explore the relative contributions of proliferation and actomyosin contractility in controlling curvature, height, and nuclear positioning.

**S-3.1 Computational model initial conditions.** Starting with the same initial tissue configuration, the quasi-steady states of perturbed model tissues are compared with the reference model tissue presented in the previous publication[9]. Comparisons were drawn at 50,000 AU (arbitrary unit) in simulation time, where each simulation time step size is 0.002 AU similar to the previous publication[44]. The midsection of the tissue can maintain its relative flatness via balancing the apical and basal contractility (Fig. 4E). Furthermore, the initial condition for simulations with cell proliferation is a curved shape representing the 72-hour AEL mark of the wing disc development (Fig. 1B-i). The default value for the spring coefficients of both apical and basal contractile springs is set as $9.0\ \mu N/\mu m$.

### S4. Image analysis and data quantification pipelines

**S-4.1 Quantification of nuclear positioning.** StarDist[10], an open-source deep learning-based ImageJ plugin was used to segment nuclei from the background (Supplementary Fig. 11A, A’). The segmentation mask was imported to MATLAB for further post-processing. MATLAB’s image labeler was first used to generate a mask defining the pouch region to separate out the nuclei of squamous epithelia from the analysis (Supplementary Fig. 11B). Fluorescence visualizing the Actin cytoskeleton was used for this task. The same image was also used to get user input of points located on the apical and basal surface using MATLAB’s *ginput()* function. Splines were next fit on the user-defined points to generate a smooth and continuous representation for the apical and basal surfaces (Supplementary Fig. 11B’). The *Regionprops* command within MATLAB was then used to estimate the centroid of each nucleus. Minimum distances between the centroid of the nuclei and the apical-basal surfaces were calculated using the *distance2curve* MATLAB function (Supplementary Fig. 11C,C’). The proximity of a nucleus from the basal surface is defined as the ratio of its distance from the basal surface over the sum of the distances from both the apical and basal surfaces. This quantity was used to generate a heatmap where each nucleus is color-coded with its proximity to the basal surface (Supplementary Fig. 11D). It can be clearly seen that as one moves towards the apical surface, the color transitions indicate an increase in the quantity.

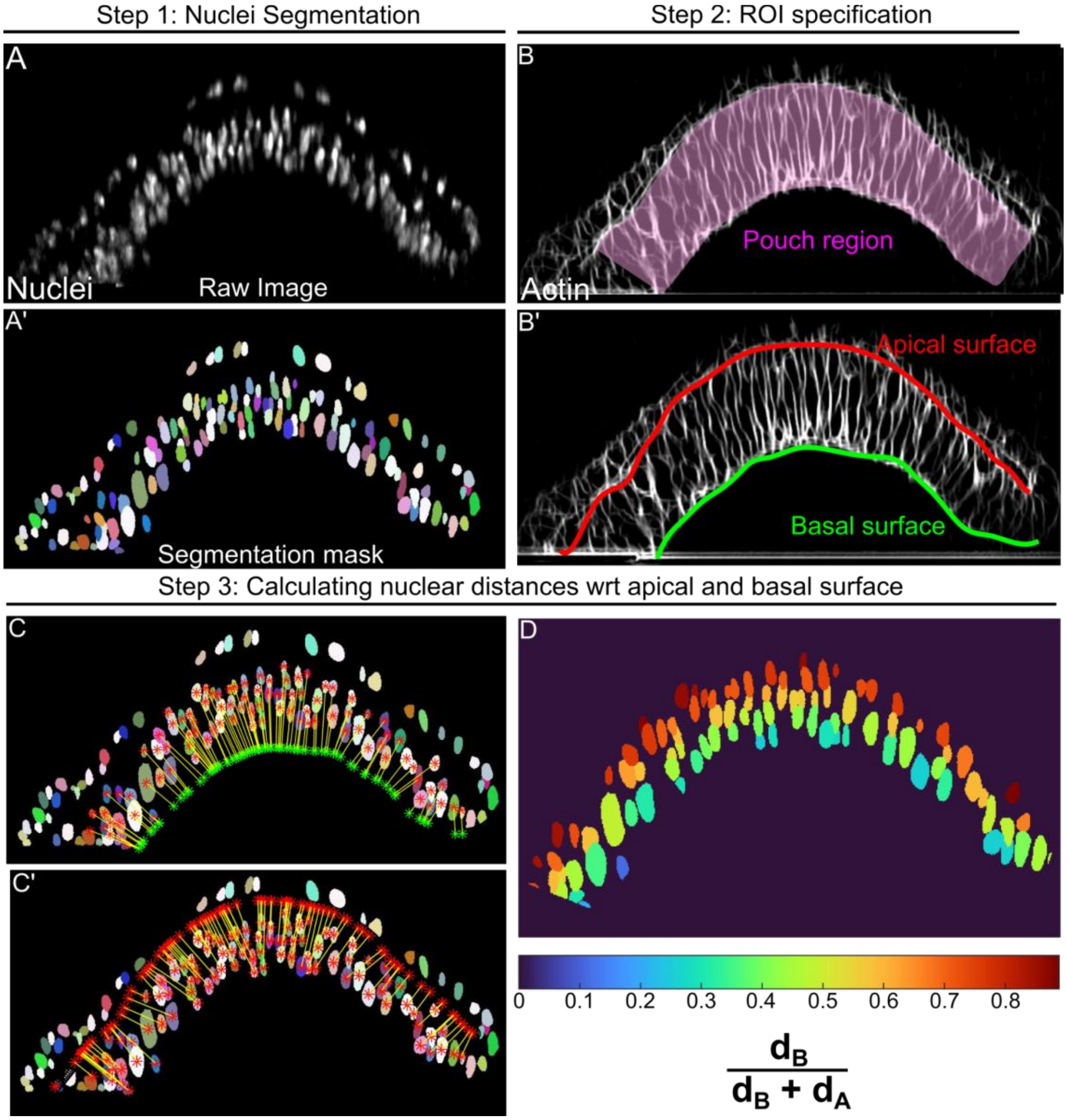


**Supplementary Figure 11. Quantification of nuclear positioning. (A)** DV cross-section of a 72-hour AEL larval wing imaginal disc with fluorescence showing nuclei. **(A')** Segmentation mask generated using StarDist. Each color represents a different nucleus. **(B)** MATLAB's image labeler was used to define the pouch region using the fluorescence channel highlighting the actin cytoskeleton. **(B')** User-defined points on the apical and basal surfaces are used to generate splines. **(C, C')** Visualization of distances of the nuclear centroids from the apical and basal surfaces respectively. **(D)** Heatmap visualizing relative positioning of nuclei with respect to the basal surface. It is defined as the distance of a nucleus from the basal surface ($d_B$) over the sum of distances of nuclei from the apical ($d_A$) and basal ($d_B$) surfaces.

**S-4.2 Quantification of morphological and signaling-related features from the wing imaginal discs.** Using Kappa, an ImageJ plugin, user-defined points were used to fit splines along the apical and basal surfaces, respectively. The tool was also used to extract fluorescent intensity of cytoskeletal regulators such as pMyoII along the points on the apical and basal surfaces. In a custom MATLAB-based tool, the number of discretization elements is first defined for analysis ($N_{cells}$). It is first used to split the basal curve into $N_{cells}$ + 1 equidistant nodes. Using the discretized points in the basal surface, corresponding points on the apical surface area are then obtained in a way such that the distances between those points and the apical surface of the pouch are minimized. The methodology allows discretization of pouches into $N_{cells}$ computational elements each mimicking a pouch cell (Supplementary Fig. 12A). A frustum-like shape obtained by joining any two consecutive points on the basal surface and the corresponding distance minimizing apical points is considered as a computational cell for our analysis. For any computational cell, the median intensity of all the points lying between the corresponding nodes on the apical and basal surface is used to approximate local pMyoII intensities across the apical and basal surfaces. Local height is defined as the average distance between the nodes on the basal surface and their counterparts on the apical surface. For a sample disc corresponding to 90 hours AEL, the pipeline was used to quantify raw intensities of pMyoII in the apical and basal nodes (Supplementary Fig. 12B). The ratio of apical and basal pMyoII intensity across each computational cell (Supplementary Fig. 12C) along with the variation of local tissue thickness (Supplementary Fig. 12D) have also been plotted.

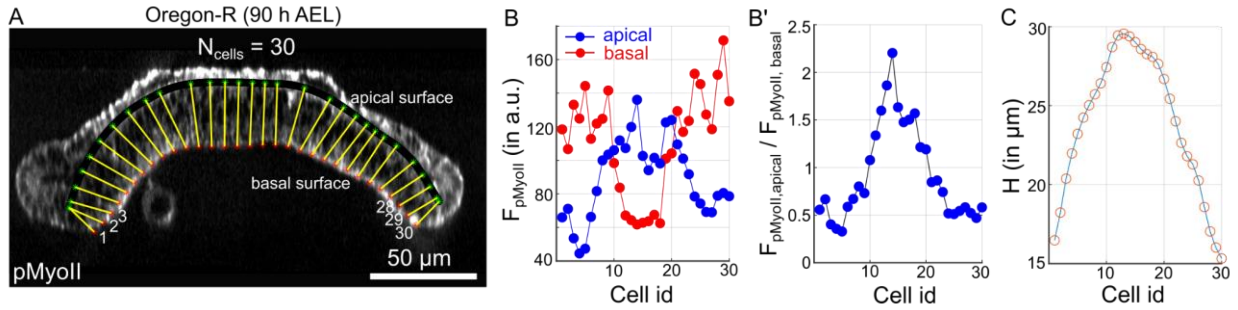


**Supplementary Figure 12. Pipeline for extracting local tissue height and the apical-basal ratio of pMyoII. (A)** Visualization of 30 discretized computation cells mimicking columnar cells within the pouch. Cell ids have been indicated in the region below the basal surface of the pouch. **(B)** Plot showing fluorescence intensity of myosin at the apical and basal surfaces of the columnar cells within the pouch. **(B')** Plot showing ratio of apical to basal pMyoII for each computational cell within pouch. **(C)** Plot showing the variation of local tissue height across the computational cells.

### S5. Supplementary Videos

**Supplementary Video 1:** (Reference to Fig. 5) Simulation showing tissue shape changes on increasing the ratio of apical to basal contractility in the medial domain of pouch. The patterning of parameters has been defined according to Fig. 5F.

**Supplementary Video 2:** (Reference to Fig. 7) Simulation showing transition in tissue shape upon increasing proliferation in only the posterior compartment (right hand side) of the wing imaginal disc. Proliferation was increased by decreasing the cell cycle length of epithelial cells in the posterior compartment by 72.5%.

### S6. Tables

| Energy & model parameters | Definition | Interaction Type | Values |
|---|---|---|---|
| $E_v$ | Volume exclusion, membrane-membrane | Morse | $U_v = -V_v = 14.08\ nN\mu m$<br>$\xi_u = 0.375\ \mu m$<br>$\xi_v = 0.094\ \mu m$ |
| | Volume exclusion, membrane-nuclei | Morse | $U_v = -V_v = 14.08\ nN\mu m$<br>$\xi_u = 0.32\ \mu m$<br>$\xi_v = 0.039\ \mu m$ |
| $E_{nuc}$ | Size of nucleus | Morse | $U_v = V_v = 35.5\ nN\mu m$<br>$\xi_u = 0.392\ \mu m$<br>$\xi_v = 5.88\ \mu m$ |
| | Interaction range | N/A | $2.1\ \mu m$ |
| $E_{ecm}$ | Volume exclusion, membrane-ecm | Morse | $U_v = -V_v = 14.08\ nN\mu m$<br>$\xi_u = 0.1375\ \mu m$<br>$\xi_v = 0.033\ \mu m$ |
| $h$ | Increment of the height of the portion of a cell with high actomyosin presence during mitotic rounding | N/A | $4.6703 \times 10^{-5} \mu m$ per simulation time step A.U. |
| Probability of in-plane cell division | The likelihood of a newly created daughter cell resides in the same cross plane of the model wing disc | N/A | 0.25 |
| Mitotic event marker | Progression (%) of cell cycle needed to enter mitosis | N/A | 95 |

**Table S1: Updated model parameters used in the subcellular element model**
